# Phylogenetic and population genetic analyses reveal patterns of divergence among isolates in the *Ceratocystis manginecans* complex

**DOI:** 10.1101/2025.08.21.671655

**Authors:** Kira M. T. Lynn, Michael J. Wingfield, Leonardo S. S. Oliveira, Acelino C. Alfenas, Rafael Ferreira Alfenas, Seonju Marincowitz, Irene Barnes

## Abstract

The taxonomic boundaries within the *Ceratocystis manginecans* complex have remained contentious due to limited morphological variation, interfertility in laboratory mating studies, and the application of different species concepts. However, recent studies have highlighted significant differences among isolates, such as host associations and biological traits, all with important implications for disease management. We examined the phylogenetic relationships, genetic diversity, and population structure of isolates treated broadly in the *Ceratocystis manginecans* complex, specifically those treated as *C. eucalypticola* and *C. manginecans* in various studies. This study included a comprehensive dataset of isolates from multiple hosts and regions, including both historical and recent outbreaks of Ceratocystis canker and wilt disease. Detailed morphological comparisons for representative isolates residing in discrete clades were conducted. Phylogenetic analyses of sequence data for seven gene regions, supported by multilocus genotyping of 16 SSR loci, identified two genetically distinct groups that broadly separated isolates having distinct ecological characteristics, the most relevant of which are infections primarily of roots or above ground tree parts. Morphological comparisons also provided support for the two groupings. Although the results supported the existence of two groups in the *C. manginecans* complex, there was also evidence of hybridization between isolates in the respective groups suggesting incomplete reproductive isolation, leading to taxonomic ambiguity. Isolates in the two groups are, therefore, treated as distinct lineages within the *C. manginecans* complex, recognizing their divergence while maintaining a conservative taxonomic approach.

## 1. Introduction

The genus *Ceratocystis* includes several economically important plant pathogens, the most aggressive of which reside in the Latin American Clade (LAC). Within the LAC, the *Ceratocystis manginecans* complex, as re-defined by Harrington et al. (2023), includes isolates that were previously considered distinct species, such as *C. manginecans*, *C. eucalypticola*, *C. mangivora*, and *C. mangicola*. These isolates are important because they cause serious diseases, particularly on trees important to plantation forestry and fruit production (Harrington et al., 2023; Liu et al., 2021). Despite causing similar disease symptoms on different hosts, which include wilting, vascular staining, bark lesions and stem cankers (Roux et al., 2020; Tarigan et al., 2011; van Wyk et al., 2012, 2007), isolates in this complex exhibit important biological and ecological differences, particularly in their host range and infection biology (Al Adawi et al., 2013b; Anjum et al., 2021; Barnes et al., 2003; Chi et al., 2019; Li et al., 2016; Liu et al., 2021; Pratama et al., 2021a, 2021b, 2024; Razzaq et al., 2020; Roux et al., 2020; Tarigan et al., 2011a; van Wyk et al., 2007). These differences are relevant because they directly affect disease management strategies including quarantine regulations, highlighting a need to accurately distinguish between isolates in the complex.

The identification and taxonomic placement of isolates in the *C. manginecans* complex has been problematic for many years. Confusion has arisen due to a number of factors including the presence of multiple ITS types in single isolates (Harrington et al., 2014; Naidoo et al., 2013), limited variation in morphology, the phylogenetic incongruencies using DNA sequences from different coding regions (de Beer et al., 2014; Fourie et al., 2015; Kanzi et al., 2020), and the fact that isolates of previously defined species are interfertile under laboratory conditions (Harrington et al., 2023). As a result, isolates have either been considered as closely related taxa, or as members of a single species complex. For example, some isolates in the *C. manginecans* complex were first described as distinct species (van Wyk et al., 2012, 2011, 2007). However, these isolates are now recognized as residing within a complex previously considered conspecific with *C. fimbriata* (Harrington et al., 2023). *Ceratocystis fimbriata* itself is now understood to be a clonal lineage and a distinct taxon (Harrington et al., 2023), even though isolates can mate in culture with those in the *C. manginecans* complex (Ferreira et al., 2010; Fourie et al., 2018; Harrington et al., 2014).

Although not primarily used for species delimitation, population genetic studies have been applied to provide insights into species or lineage boundaries, especially for species complexes (Choi, 2016; Hey and Pinho, 2012). Such studies assess genetic diversity and structure by analysing allele frequencies, genetic drift, gene flow, mutation, and selection. Previous population genetic studies on isolates in the *Ceratocystis manginecans* complex (formerly designated as *C. manginecans* and *C. eucalypticola*) have focused on aspects such as genetic diversity, pathways of movement, host associations, and possible origins of these economically important fungi (Al Adawi et al., 2014; Ferreira et al., 2013; Fourie et al., 2016; Hlongwane, 2022; Li et al., 2016; Liu et al., 2021). However, confusion regarding their taxonomic placement has hindered comparative studies. For example, Fourie et al. (2016) suggested that Southeast Asia (SEA) could be the origin of isolates previously identified as *C. manginecans*. In contrast, studies in Brazil revealed significant genetic variation, implying that the pathogen (previously designated as *C. fimbriata*) is native to Brazil and has spread globally through infected *Eucalyptus* propagation material (Ferreira et al., 2013, 2011; Harrington et al., 2023; Li et al., 2016). Liu et al. (2021) compared isolates from SEA and Brazil but could not fully resolve lineage boundaries due to limited representation of Brazilian isolates (Harrington et al., 2023; Oliveira et al., 2015). However, their study did reveal two major genetic clusters based on host: one from *Eucalyptus* spp. (previously designated as *C. eucalypticola*) and another from *Acacia* spp. (designated as *C. manginecans*). Similar host association patterns were reported by Fernandes et al. (2024), suggesting ongoing speciation driven by host associations and multiple host expansion events.

Subsequent to the first description of *C. manginecans* on *A. mangium* by Tarigan et al. (2011), serious outbreaks of disease caused by *Ceratocystis* spp., particularly on *Acacia* spp. and *Eucalyptus* spp., have continued to occur in SEA. These include outbreaks in Indonesia (Hlongwane, 2022; Pratama et al., 2021a, 2021b, 2024; Suwandi et al., 2021), Malaysia (Wingfield et al., 2023), and Vietnam (Thu et al., 2024), There have also been outbreaks of the disease on *Eucalyptus* in South Africa (Hlongwane, 2022; Roux et al., 2020) and Brazil (Benso et al., 2024; Ferreira et al., 2011). In addition to those collected from diseased trees, isolates of *Ceratocystis* spp. have also been collected from the newly exposed surfaces of recently felled *Eucalyptus* in China (Liu et al., 2021) and more recently in Colombia (C.A. Rodas, unpublished).

Recent outbreaks of disease caused by isolates residing in the *C. manginecans* complex as defined by Harrington et al. (2023), coupled with evidence of relevant differences in host range, geographic distribution and biology of isolates (Fernandes et al., 2024; Liu et al., 2021), have underscored a need to explore how these differences are reflected in genetic clustering. Following this reasoning, the objective of our study was to facilitate the accurate characterization and identification of isolates within the complex, linking them to distinct genetic clusters and traits, important for disease management and quarantine. This was achieved by studying an extensive dataset for *Ceratocystis* isolates previously identified as *C. manginecans* or *C. eucalypticola,* from both historical and recent outbreaks, and considering their multilocus phylogeny, genetic diversity and population structure.

## 2. Materials and methods

### 2.1 Fungal isolations and DNA extractions from new locations

Samples were collected specifically for this study from countries experiencing new outbreaks of *Ceratocystis* infections on trees, including Brazil, South Africa, Malaysia and Indonesia (Supplementary Table 1). These samples were from either new host species or newly affected regions, with collections carried out randomly across sites (Supplementary Table 1). Common symptoms of infection included wilting and brown-streaked discolouration of the sapwood. Isolates from Colombia were collected from the stumps of freshly harvested *Eucalyptus* trees.

Isolations were made by placing ∼50 g of thin shavings from discoloured wood between two 1 cm carrot discs soaked overnight in ddH₂O amended with 0.001 g/vol streptomycin sulphate (SIGMA, Steinheim, Germany). These carrot baits were placed in moist chambers and incubated at 28 °C for up to two weeks to stimulate development of *Ceratocystis* ascomata (Moller and De Vay, 1968). Alternatively, wood pieces were incubated in sealed plastic bags to maintain moisture and induce sporulation. Where present, single ascospore drops were lifted from ascomata and transferred to Petri dishes with 2 % malt extract agar (MEA: 20 g/L malt extract from Biolab, 20 g/L agar from Difco), incubated 10 days at 25 °C. Pure cultures on 2 % MEA were produced using single hyphal tip transfers. Only one isolate per tree was selected for downstream processing.

After 10 days at 25 °C, mycelium was scraped from agar surfaces with a sterile scalpel and genomic DNA extracted using the Zymo Quick-DNA Fungal/Bacterial Kit per manufacturer’s instructions. DNA was standardised to 30 ng/µl. Selected isolates from these collections were used in phylogenetic analysis, and all isolates were SSR genotyped as described below. All isolates are maintained in the Forestry and Agricultural Biotechnology Institute (FABI) culture collection (CMW), Pretoria, South Africa.

### 2.2 DNA and data acquisition from other studies

To combine previously generated data with new isolates, all available ITS, MS204, and *rpb2* sequences, and microsatellite allele data, were obtained from De Mar Angel (unpublished), Hlongwane (2022), Fourie et al. (2016), Liu et al. (2021) and Lynn (2017). Data from Liu et al. (2021) and Hlongwane (2022) included only 10 SSR markers, thus DNA from those studies were retrieved to generate missing SSR data as described below. DNA from isolates obtained from diseased *Eucalyptus* in Brazil, for which no prior molecular data existed, was also included.

### 2.3 PCR amplification, sequencing and phylogenetic analysis

All newly acquired DNA samples were sequenced to confirm species identity. The internal transcribed spacer (ITS) rDNA region and 5.8S rRNA gene were amplified with primers ITS1 and ITS4 (White et al., 1990). The guanine nucleotide-binding protein subunit beta-like protein (MS204) gene was amplified with primers MS204F.ceratoB and MS204R.ceratoB (Fourie et al., 2015) to further confirm species identity. PCR reactions for both loci contained of 2.5 μl 5× MyTaq buffer (Bioline, London, UK), 0.25 μl MyTaq DNA polymerases (Bioline), 1 μl DNA template, 0.5 μl of each primer (10 mM), and 8.25 μl of sterile deionized water, for a 13 μl total reaction mixture. The PCR cycler program was 95 °C for 5 min, 10 cycles of 95 °C for 30 s, 56 °C for 45 s, 72 °C for 90 s, another 30 cycles of 95 °C for 30 s, 56 °C for 45 s and 72°C for 90 s (5 s increase per cycle at 72 °C) with a final step at 72 °C for 10 min. PCR amplification success was evaluated using agarose gel electrophoresis (AGE). Amplicons were purified using ExoSAP-IT PCR Product Clean-up Reagent (Thermo Fisher Scientific), and sequenced bidirectionally using BigDye Terminator v3.1 (Applied Biosystems, Forster City, California) with the same primers. The thermal cycling conditions included 25 cycles of 10 s at 96 °C, 5 s at 56 °C, and 4 min at 60 °C. Sequencing was run on an ABI PRISM 3100 (Applied Biosystems) at the University of Pretoria.

Forward and reverse sequencing reads were assembled into contigs using CLC Bio Main Workbench 6 (CLC Bio, www.clcbio.com), and the consensus sequences generated were exported for phylogenetic analyses. A preliminary identity for the isolates was obtained by performing a nucleotide BLAST search of the ITS and MS204 sequences against the National Centre for Biotechnology Information (NCBI) GenBank database (http://www.ncbi.nlm.nih.gov), using the ‘type’ only setting. Based on the results, the sequences generated in this study, and those designated as *C. manginecans* in Harrington et al. (2023), were incorporated into datasets of Latin American Clade (LAC) ex-type species (Barnes et al., 2018; Supplementary Table 2). Alignments were made in MEGA v7 (Kumar et al., 2016), with MUSCLE alignment software (Edgar, 2004). Resulting alignments were used to construct separate phylogenetic trees of the ITS and MS204 gene regions based on Maximum likelihood (ML), using raxmlGUI 2.0 (Edler et al., 2021). A non-parametric analysis of the sequence data with 1000 bootstrap replicates provided statistical support for the branches of the generated ML trees. *Ceratocystis albifundus* CMW4068 was used to root the ML trees.

Based on the initial sequence identification of the isolates and the topology of the resulting phylogenetic trees, representative isolates from each of the major clades were selected for additional DNA sequence analyses to confirm the identity of isolates (Supplementary Table 1). This included sequencing the β-tubulin 1 (βT 1) region using the primers βT1a and βT1b (Glass and Donaldson, 1995), the Transcription Elongation Factor-1 alpha (tef1) gene region with primers TEF1F and TEF2R (Jacobs et al., 2004), and the second largest subunits of RNA polymerase II (rpb2) using the primers RPB2-5Fb and RPB2-7Rb (Fourie et al., 2015). PCR and sequencing reactions were performed as described above with locus-specific annealing temperatures (Fourie et al., 2015). Resulting sequences for the additional gene regions were assembled and used to construct phylogenetic trees using the same protocols described above. The four individual data sets for the MS204, βT 1, tef1 and rpb2 regions were combined using FASconCAT-G (Kück and Longo, 2014) and a final combined multi-gene ML phylogenetic tree was generated using the same parameters stated above. *Ceratocystis albifundus* CMW4068 was used to root the ML tree.

Several representative isolates from each clade identified based on the topology of the ITS and MS204 ML trees, were also sequenced for MAT 1 (*MAT1-1-2*) and MAT2 (*MAT1-2-1*) regions. The MAT gene regions were amplified to screen isolates used in this study against the five newly described LAC species described by Harrington et al. (2023), as the gene regions used in this study were not available for those species. The MAT1 and MAT2 regions were amplified and sequenced with the primers CFMAT1-F and CFMAT1-R, and X9978R1R and CFM2-1F respectively, following the PCR and sequencing cycling reactions described by Harrington et al. (2014). The resulting sequences were incorporated into datasets of the LAC ex-type species (Harrington et al., 2023; Supplementary Table 2) and used to construct individual phylogenetic trees using the same protocols described above. *Ceratocystis albifundus* C1060 (CMW2475) was used to root the ML tree.

### 2.4 Microsatellite marker genotyping

Sixteen microsatellite markers previously shown to be informative for *Ceratocystis* (Barnes et al., 2001; Fourie et al., 2016; Steimel et al., 2004), were used to genotype all isolates (Supplementary Table 1). PCR mixtures and cycling programs followed published protocols (Barnes et al., 2001; Fourie et al., 2016; Steimel et al., 2004). Successful amplification was verified with AGE.

Amplicons were run on GeneScan (Applied Biosystems, Thermo Fisher Scientific, Carlsbad, USA), using panels described by Fourie et al. (2016). Amplicons for each respective panel were pooled together in a 1/200 dilution mixture with sterile water, from which 1 μl of the resulting mixture was combined with 0.2 μl Liz500(−250) size standard (Applied Biosystems, Thermo Fisher Scientific) and 10 μl formamide, and run on an ABI PRISM^TM^ 3500xl Auto sequencer at the University of Pretoria (Thermo Fisher Scientific, Carlsbad, CA, USA). Fragment sizes were scored with GeneMapper® v. 6 software (Applied Biosystems, Thermo Fisher Scientific). Multi-locus genotypes (MLGs) were generated by combining alleles across the 16 loci. Markers missing from SSR data from previous studies (Hlongwane, 2022; Lui et al., 2021) were amplified and scored as described above and incorporated into the larger dataset.

Several representative alleles, including unique alleles when present, for each of the 16 loci investigated in this study, were selected for sanger sequencing to confirm the allele size obtained with GeneScan. These alleles were amplified and sequenced with unlabelled microsatellite primers following the protocols described by Barnes et al. (2001), Fourie et al. (2016) and Steimel et al. (2004). Sequences were assembled into contigs and manually inspected using CLC Bio Main Workbench v. 6 (CLC Bio, www.clcbio.com).

### 2.5 Population genetic analyses

#### 2.5.1 Genetic diversity statistics

To calculate various diversity statistics, data for all isolates (newly acquired and those from previous studies) were grouped into populations based on their country of origin, regardless of their previous identification as *C. manginecans* or *C. eucalypticola.* This approach allowed investigation of whether any genetic clustering would align with the lineages in the *C. manginecans* complex, as determined through multi-gene phylogenetic analyses. Additional diversity statistics were calculated for sub-populations identified within these lineages. Analyses used the R package *poppr* (Kamvar et al., 2014) in R v. 4.0.0 (R Core Team, 2020) run on RStudio (RStudio Team, 2020), on the non-clone-corrected dataset. Statistics included total MLGs, expected MLGs (eMLG; Hurlbert, 1971), genetic diversity (Hexp: Nei, 1978), Shannon-Wiener index (H: Shannon, 2001), Stoddart and Taylor’s index (G: Stoddart and Taylor, 1988), Simpson’s index (λ: Simpson, 1949), and genotypic evenness (E_5_: Grünwald et al., 2003).

#### 2.5.2 Population structure

The Bayesian clustering algorithm in STRUCTURE 2.3.4 (Falush et al., 2003), was used to assign individuals to clusters (K) and infer population structure. Clone-correction was carried out on the dataset per country using the *clonecorrect()* function in R. The clone-corrected dataset per country was analysed, testing K = 1–20 with 20 runs per K and using admixture and independent allele frequency models (Pritchard et al., 2000). The first 10 000 Markov Chain Monte Carlo (MCMC) iterations of each run were discarded as burn-in, and 100 000 MCMC iterations were retained for analysis. The optimal K was determined using StructureSelector (Li and Liu, 2018), implementing the Evanno method (Evanno et al., 2005). After the optimal K was determined a final STRUCTURE analysis was performed, testing K = 1 to 10, in 20 independent runs for each K where the first 100 000 MCMC iterations of each run was discarded as burn-in, and 1 000 000 MCMC iterations were retained for analysis. CLUMPAK (Kopelman et al., 2015) incorporated into StructureSelector was used to merge runs and visualise results. Because STRUCTURE detects major variation in the dataset (Evanno et al., 2005), additional analyses were run on non-clone-corrected subsets from each cluster in the initial run, guided by the multi-gene ML results, using the same parameters.

Population genetic structure was also visualised with discriminant analysis of principal components (DAPC; Jombart et al., 2010) in R v. 4.0.0 (R Core Team, 2020) run on RStudio (RStudio Team, 2020). DAPC was conducted using the *adegenet* package (Jombart and Ahmed, 2011) on non-clone-corrected data. The optimal cluster number was selected via Bayesian information criterion (BIC), and the *xvalDapc* function was used to determine the number of principal components to retain in the analysis (PCA). DAPC analyses were also performed on isolates showing mixed ancestry to assess hybrid zones.

### 2.6 Minimum spanning network analyses

Relationships among individuals were visualised with a minimum spanning network (MSN) using Edwards (1971) genetic distances with the R packages *poppr* (Kamvar et al., 2014) and *ape* (Paradis et al., 2004). Edwards genetic distances were selected for this analysis as it effectively captures genetic variation among isolates, making it suitable to resolve species boundaries and population structure for the extensive dataset analysed in this study. An MSN was also generated for isolates showing mixed ancestry.

### 2.7 Calculation of pairwise population differentiation and gene flow

Analysis of molecular variance (AMOVA: Excoffier et al., 1992), was conducted to determine whether genetic variation is differentiated within and among the populations. AMOVA was performed in R v. 4.0.0 run on RStudio (RStudio Team, 2020) using the package *poppr* (Kamvar et al., 2014), with the *poppr.amova* function on the non-clone-corrected dataset. Significance was tested using 1000 permutations where the null hypothesis of no genetic differentiation was rejected at p ≤ 0.001. AMOVA was also performed for mixed-ancestry isolates.

Pairwise population differentiation (ϕPT) was calculated using Hedrick’s standardised Gst (Hedrick, 2005) with the *pairwise_Gst_Hedrick* function in *poppr* (Kamvar et al., 2014) for the non-clone-corrected dataset. Gene flow (Nm) was calculated from ϕPT (Slatkin and Barton, 1989). Analyses were conducted for populations defined by country and for the two ML tree lineages. Gst values near 1 indicate high differentiation and low gene flow, whereas values near 0 indicate low differentiation and higher gene flow. The significance of observed differentiation values was tested using the perm_test function in R v4.0.0 with 1000 permutations, applying significance thresholds of 0.05 and 0.01.

### 2.8 Morphological comparisons

Three *Ceratocystis* isolates from *Eucalyptus* plantations in KwaZulu-Natal, South Africa (CMW56740, CMW58001, CMW57998), and three from infected *Eucalyptus* in Riau, Indonesia (CMW60227, CMW60230, CMW60225), were selected for morphological comparison. These represented the two major clusters from the genetic analyses. The South African isolates were associated with root disease whereas the Indonesian isolates were associated with above-ground infections.

Three replicates per culture per phylogenetic group (18 total) were randomized and assessed blindly. Replicates were grown on 60 mm Petri dishes containing 2 % MEA in the dark at 25 °C for seven days. On the seventh day, 10–15 ascomata and their adjacent conidiophores were harvested from six equal sectors per replicate plate. Fungal structures were initially mounted in water, which was then replaced with 85% lactic acid for observation and preservation. Twenty-eight morphological characters were assessed for the comparisons (Table 4). For twenty-six characters, 25 measurements per replicate were made (225 per character). The number of unbranched or branched aleuriospore-conidiophores associated with a single ascoma was counted, and the ratio (branched: unbranched) was used in the analysis. For branching pattern, 30 ascomata per replicate were examined (270 ratio data points) Measurements were made using a Nikon microscope (Eclipse Ni, Japan) mounted with a camera (Nikon DS-Ri2) and imaging software (NIS-Elements BR, Nikon, Japan). After which data were regrouped by isolate, then phylogenetic cluster for statistical analyses. Three characters linked to the conidiogenous cells of barrel-shaped conidia were excluded from the analyses due to an absence of data in one isolate (CMW60227). Statistical analyses (α = 0.05) on 24 characters, including the ratio, were run in R (R Core Team, 2023). Outliers were detected with boxplots and Rosner’s test (Millard, 2013). Normality and homogeneity of variance were tested with Shapiro–Wilk, Bartlett’s, and Levene’s tests. Two-sample t-tests and Wilcoxon rank-sum tests assessed differences between clusters.

## 3 Results

### 3.1 Fungal isolates and DNA extractions from new locations

A total of 364 isolates morphologically typical of *Ceratocystis* were obtained from Brazil, Colombia, South Africa, Malaysia, and Indonesia (Supplementary Table 1). Most were from diseased *Eucalyptus* spp., with a small number were from diseased *Acacia* spp. in Indonesia, along with two isolates from *Lansium domesticum* and *Mimusops elengi* (Supplementary Table 1). Isolates from Southeast Asia were collected from hosts showing only above-ground infections, whereas those from South Africa were from trees with root disease. Colombian isolates were recovered from stumps of freshly harvested *Eucalyptus* in the absence of visible disease symptoms.

### 3.2 DNA and data acquisitions

DNA from 707 isolates was acquired from previous studies and collaborators (Supplementary Table 1). In total, 1192 isolates spanning 11 regions and eight hosts were analysed (Fig. 1; Supplementary Table 1).

**Fig. 1.**
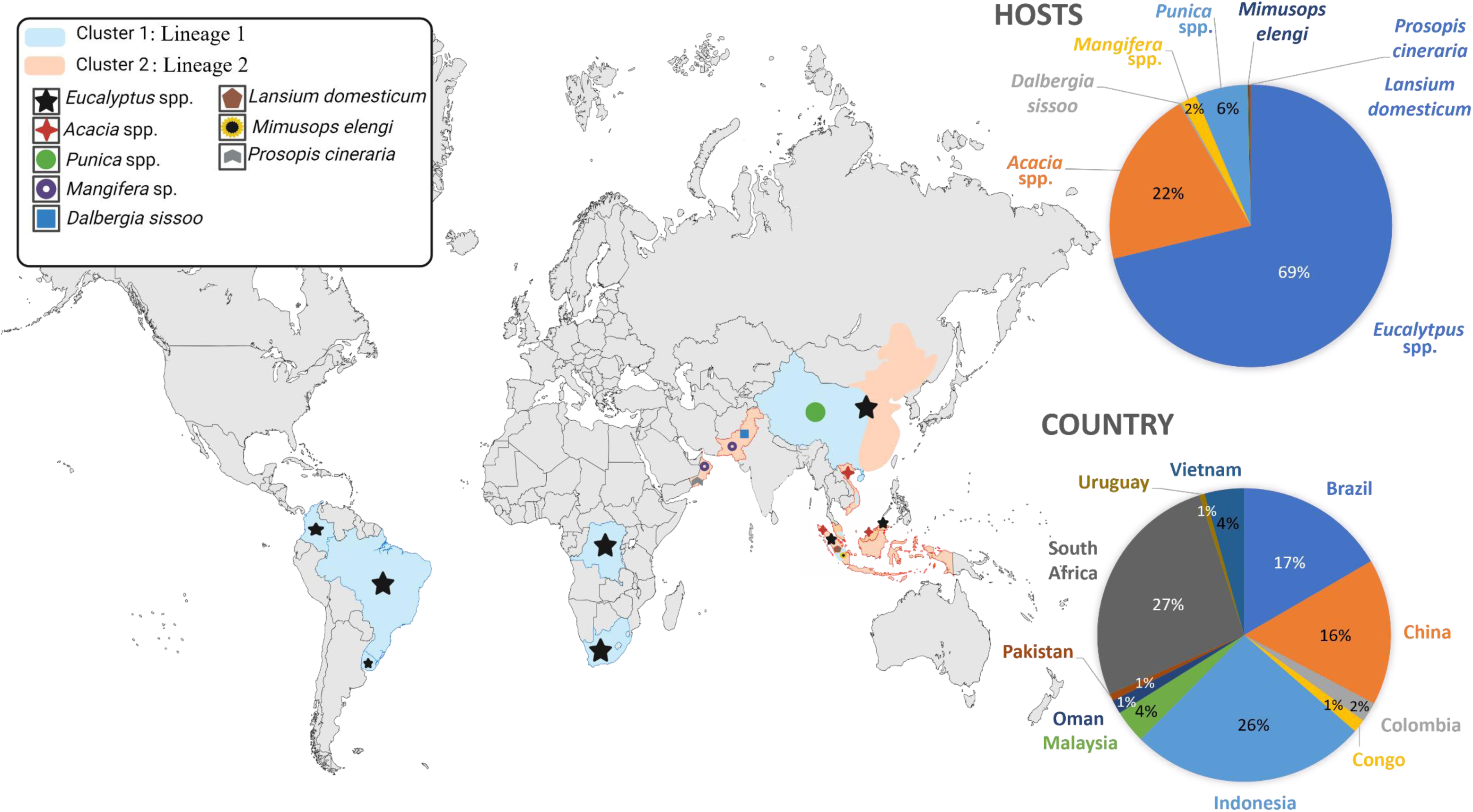
Geographical distribution of *Ceratocystis* clusters and host proportions. Distribution of *Ceratocystis* clusters across different geographical regions, including plant hosts and their proportions by country. The map illustrates the locations of clusters and their associated hosts, highlighting the relative abundance of each host species in the represented countries

### 3.3 PCR amplification, sequencing, and phylogenetic analysis

#### 3.3.1 ITS and MS204 gene regions

ITS (600 bp) and MS204 (930 bp) sequences were obtained for most newly screened isolates. ITS chromatograms for 213 isolates (Supplementary Table 1) showed conflicting sequence data due to multiple ITS types present (Naidoo et al., 2013). These ITS data were not used for identification. Based on the nucleotide BLAST results for the ITS region, 130 resided in the *C. manginecans* complex and grouped into either Lineage 1 (formerly *C. eucalypticola*) or Lineage 2 (formerly *C. manginecans*). (Supplementary Table 1). For the MS204 region, 362 isolates were similarly identified and grouped into either Lineage 1 or 2 of the *C. manginecans* complex (Supplementary Table 1). ML phylogenetic analyses of the ITS (Supplementary Fig. 1) and MS204 (Supplementary Fig. 2), supported the groping of isolates into the two lineages.

Nine ITS sequence variants were detected overall, with isolates from Brazil showing considerable variation. Variants included ITS5 (*C. eucalypticola*), ITS6 and ITS7b (*C. manginecans* ITS type 1 and 2; Al Adawi et al., 2013; Harrington et al., 2014), and six additional variants. Two Brazilian isolates (Variant 4) formed a monophyletic in the ITS phylogeny with high statistical support (74 %; Supplementary Fig. 1: red). Several other isolates (all from Brazil) displayed fixed SNP variations across multiple isolates, forming three additional ITS sequence variants (Variant 5–7). Statistical support for these SNP variations was low and clustered these variants together with *C. cacaofunesta* (CMW14798) and *C. manginecans* (C1442) (Supplementary Fig. 1: black). Similarly, some Colombian isolates showed fixed SNP variations, forming two additional ITS sequence variants (Variant 8 and 9), with low statistical support (Supplementary Fig. 1: yellow). Due to the low bootstrap support and the known limitations of the ITS gene region for this genus, ITS was excluded from combined analyses.

#### 3.3.2 tef, βT 1 and rpb2 gene regions

Across the representatives of the newly acquired isolates screened, the DNA sequence lengths were 725 bp for tef, 580 bp for βT 1 and 1130 bp for rpb2 loci. Maximum likelihood phylogenetic analyses of the tef1 (Supplementary Fig.3), βT 1 (Supplementary Fig. 4) and rpb2 (Supplementary Fig. 5) further confirmed the identity of all isolates, grouping them within the *C. manginecans* complex. ML phylogenetic analyses of the tef1 grouped all but five representatives of the newly acquired isolates together with the ex-type species of *C. eucalypticola, C. manginecans*, *C. cacaofunesta, C. mangivora, C. fimbriata* and *C. fimbriatomima* (Supplementary Fig. 4) as this gene region does not distinguish between these lineages (Fourie et al., 2015). The remaining five isolates, all from Brazil, formed a monophyletic clade with high statistical support (75 %), most closely associated with the above grouping (Supplementary Fig. 4: green). Likewise, ML phylogenetic analyses of the βT 1, grouped all representatives of the newly acquired isolates together with the ex-type species of *C. eucalypticola, C. manginecans* and *C. curvata* (Supplementary Fig. 4; Fourie et al., 2015). Only the rpb2 gene region provided results consistent with the MS204 region, delineating isolates into two major lineages within the *C. manginecans* complex. Isolates from Brazil, South Africa, Colombia, and several isolates from Indonesia grouped within Lineage 1 (previously identified as *C. eucalypticola*; Supplementary Fig. 5). Isolates from Indonesia and Malaysia, along with ex-type isolates previously identified as *C. manginecans*, *C. mangicola*, and *C. mangivora*, grouped within Lineage 2 (Supplementary Fig. 5). Clustering across single-locus trees was generally congruent overall (Supplementary Fig. 2 - 5).

A combined four locus ML analysis (MS204, tef, βT1, and rpb2, with a database alignment of 3239 bp) assigned 55 newly acquired isolates to Lineage 1 (formerly designated as *C. eucalypticola*) or 2 (formerly designated as *C. manginecans*) within the *C. manginecans* complex (Fig. 2). Isolates from Brazil, Colombia, South Africa, and several from Indonesia formed a statistically supported monophyletic clade (69%), identified here as Lineage 1 (Fig. 2). Isolates from Malaysia and Indonesia formed a statistically supported monophyletic clade (74%), identified here as Lineage 2. The phylogenies of MS204 and rpb2 were the most informative gene regions, capable of delineating isolates within the *C. manginecans* complex. As a result, a combined ML phylogenetic tree (MS204 and rpb2) is presented in Fig. 3. Previously published isolates (De Mar Angel, unpublished; Fourie et al., 2016; Hlongwane, 2022; Liu et al., 2021; Lynn, 2017) were re-assigned to lineages based on this ML analysis (Fig 3).

**Fig. 2.**
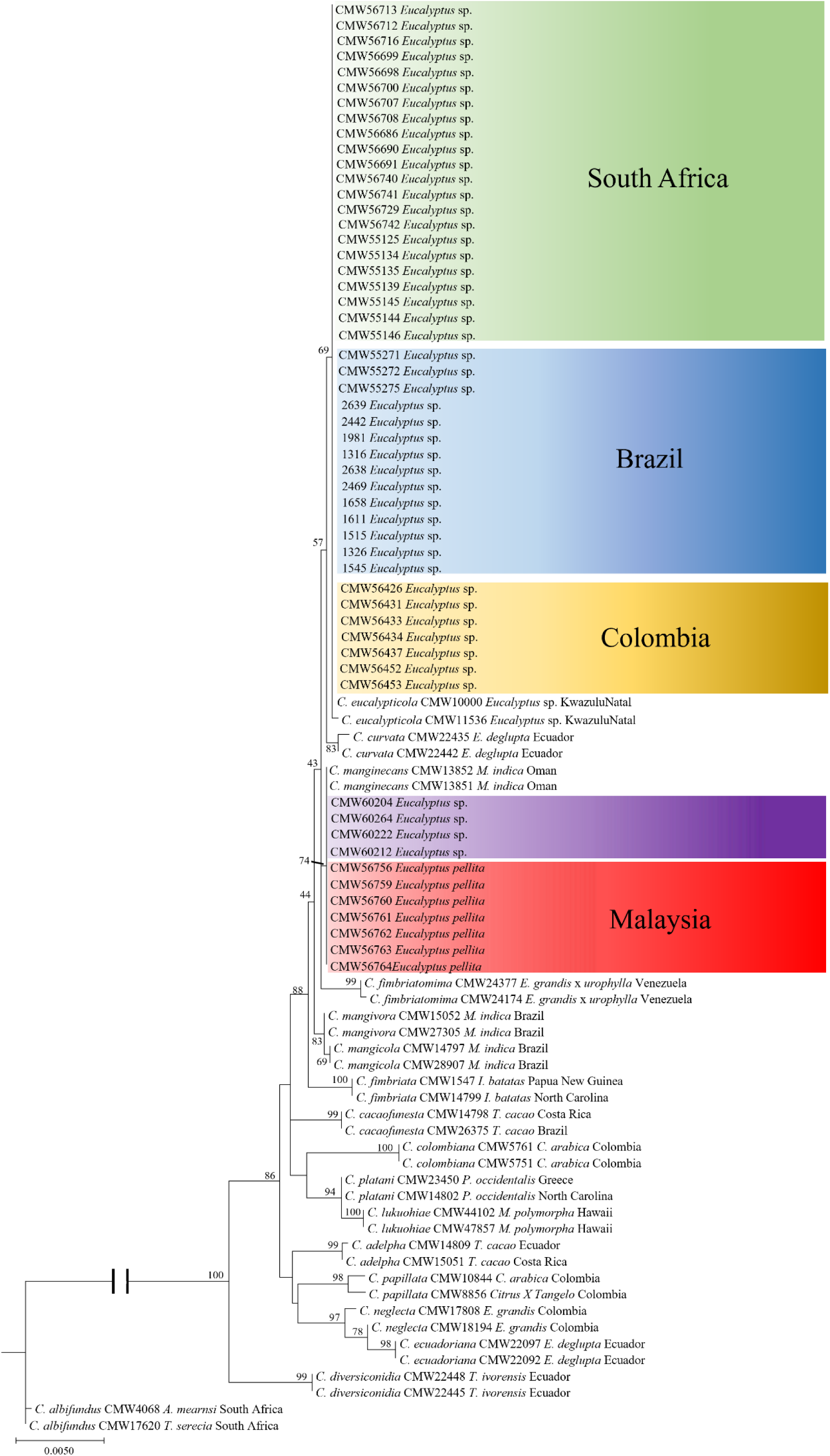
Phylogenetic tree based on multi gene maximum likelihood (ML) analysis of MS204, βT 1, tef1, and rpb2 sequences for *Ceratocystis* species in the LAC. LAC and *Ceratocystis* isolates used in this study (only representative haplotypes per country and host were included in the analysis). Coloured boxes indicate representatives of isolates sequenced in this study from five regions (Brazil, Colombia, South Africa, Malaysia, and Indonesia). Bootstrap values higher than 50% are indicated in the figure.

**Fig. 3.**
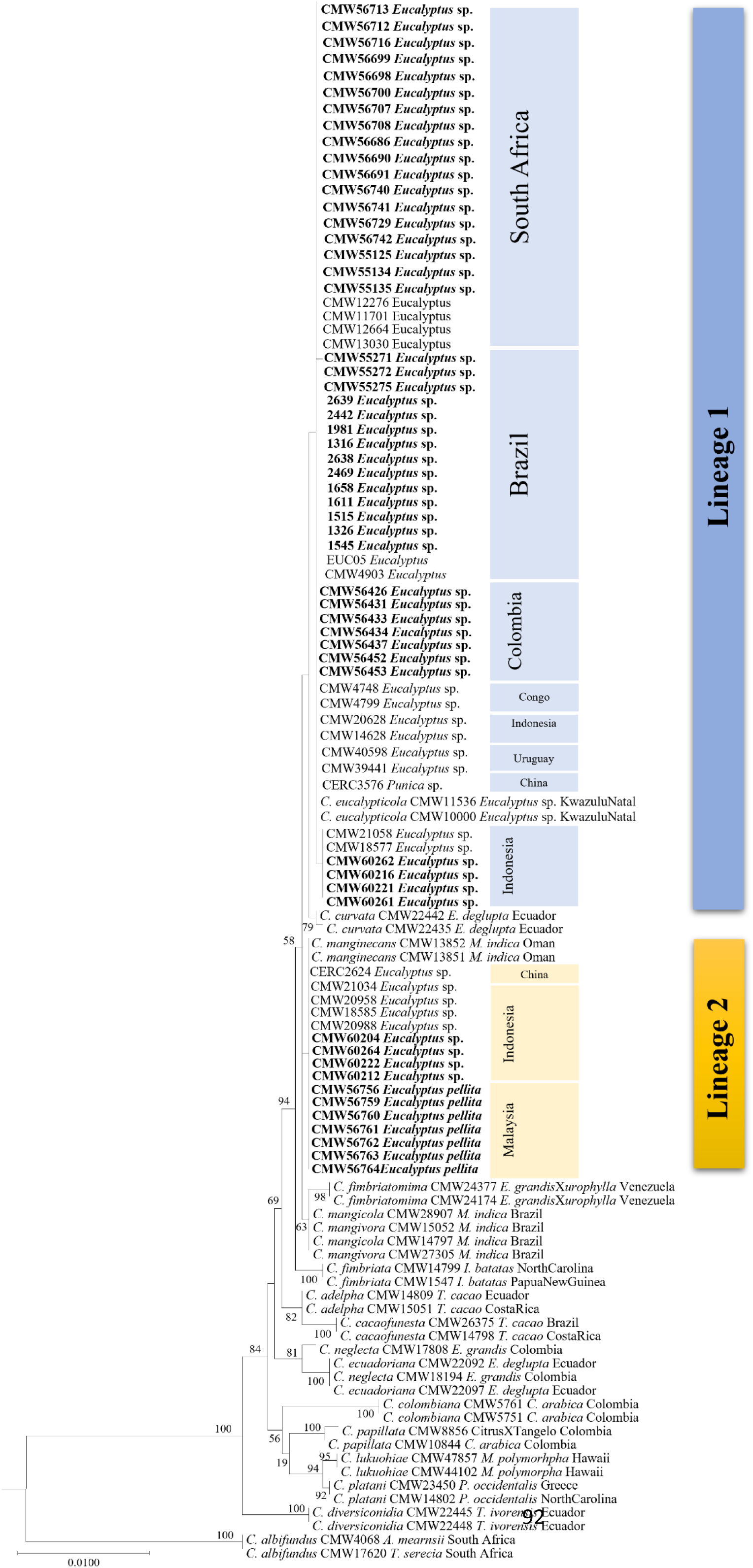
Phylogenetic tree based on maximum likelihood (ML) analysis of a combined dataset of MS204 and rpb2 gene sequences for *Ceratocystis* isolates used in this study (only representative haplotypes per country and host were included in the analysis). Isolates in bold and highlighted in coloured blocks are the isolates sequenced in this study and were either identified as Lineage 1 or Lineage 2 in the *C. manginecans* complex. Bootstrap values higher than 50% are indicated in the figure.

Topological discrepancies occurred in 56 isolates from *Eucalyptus* in Colombia (8), South Africa (34), Indonesia (10), and Brazil (4), primarily between ITS and MS204 (49 isolates; Supplementary Table 1: Highlighted in blue). Seven Indonesian isolates showed discrepancies between MS204 and *rpb2* (Fig. 3: Black box). These patterns are consistent with mixed ancestry (hybridisation) between the two lineages.

#### 3.3.3 MAT 1 and MAT 2 gene regions

Approximately 1,130 bp were generated for each MAT locus across 96 representatives. MAT phylogenies were broadly concordant with the multi-gene tree, each recovering three well-supported clades (Supplementary Figs. 6, 7), although several isolates clustered independently as noted below.

In the MAT1 ML phylogeny, isolates formed three clades (Supplementary Fig. 6). Group 1 (97 % bootstrap support) was predominantly from Brazil, South Africa, Colombia, and Indonesia. Group 2 (97 % bootstrap support) consisted mainly of Malaysian and Indonesian isolates. Group 3 (98 % bootstrap support) comprised mostly Brazilian isolates. The remaining isolates from Brazil grouped closely with those in group 1 and a single isolate from Colombia (CMW 56431) clustered on its own but was closely related to *C. fimbriatomima*.

The MAT2 ML phylogeny also resolved three clades (Supplementary Fig. 7). Group 1 (94 % bootstrap support) contained most isolates except those from Malaysia. Group 2 (77 % bootstrap support) comprised Malaysian and Indonesian isolates. Group 3 (93 % bootstrap support) included four Indonesian isolates, clustering closest to Group 1. Again, CMW 56431 formed its own branch, closely related to *C. fimbriata* (strain C1476) and *C. fimbriatomima*.

### 3.4 Microsatellite marker genotyping

All 16 microsatellite markers were polymorphic with the number of alleles per locus ranging from three to 32. A total of 134 different alleles were generated. A total of 159 representative alleles were sequenced. Occasionally, the sequence lengths differed from that obtained with fragment analyses. Sequenced allele sizes were used to recalibrate fragment sizes for all downstream analyses (Supplementary Table 1).

### 3.5 Population genetic analyses

#### 3.5.1 Genetic diversity statistics

From 1174 isolates, 466 MLGs were detected. Genotypic diversity (G and H) varied across 11 country-level populations, with relatively high variation in six countries (Table 1). Six countries had high Simpson indices (λ > 0.90), whereas others had lower λ, suggesting a more clonal population (e.g., Oman λ = 0.20; Pakistan λ = 0.37) or a bottleneck following a recent introduction (e.g., Colombia λ = 0.51). Evenness (E5) ranged from 0.45 to 0.84, indicating a moderate to high evenness in genotype distribution (Table 1). Gene diversity (Hexp) also varied across countries, being highest in Vietnam (0.51), followed by Indonesia (0.46), Brazil (0.36), and China (0.35). The lowest Hexp values (< 0.05) were observed in Oman, Pakistan, and Colombia. Results were similar when analysed by lineage (1 and 2), although small sample sizes for some populations limited calculation of meaningful diversity statistics (Table 1). Notably, Hexp was highest in Brazil (0.36) for cluster 1 (Lineage 1) and in Vietnam (0.42) for cluster 2 (Lineage 2). In the hybrid sub-population, 31 MLGs were detected among 56 isolates, with the highest diversity in Brazil (0.24), followed by South Africa (0.16), Indonesia (0.13), and Colombia (0.03) (Table 1).

**Table 1.**
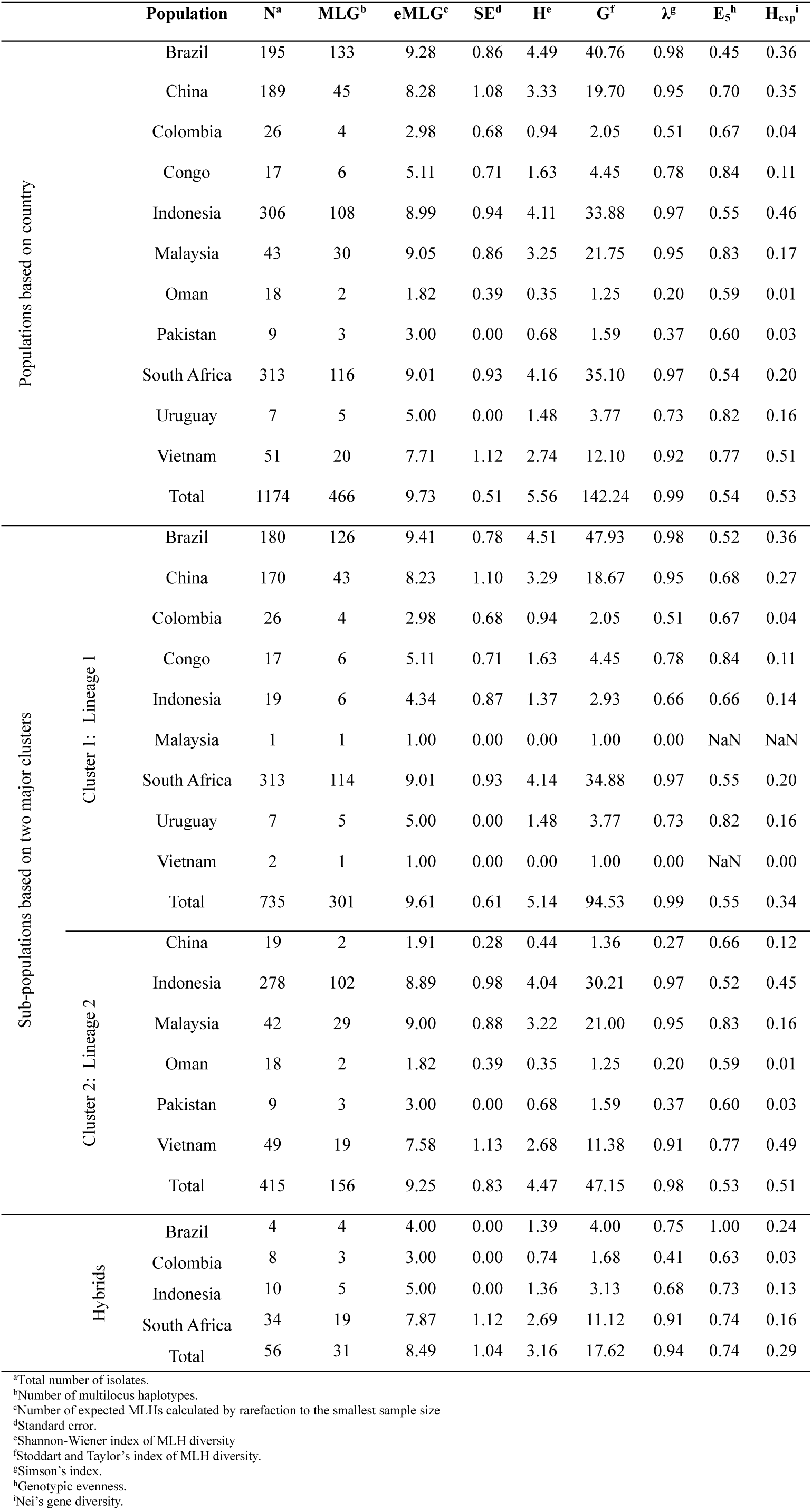
Genetic diversity statistics for isolates grouped into populations based on geographical location and on mutli-gene ML phylogenetic clusters.

#### 3.5.2 Population structure

STRUCTURE analysis without prior assumptions, indicated optimal K = 2 and K = 5 (Evanno ΔK, LnP(K); Fig. 4). For K= 2, isolates from Brazil, China, Colombia, Congo, South Africa, and Uruguay clustered together (blue) and isolates from Indonesia, Malaysia, Oman, Pakistan and Vietnam clustered together (orange). These results are congruent with the lineages observed in the multi-gene ML tree and suggested some level of geographic segregation of the clusters. Evidence of admixture was observed in several populations, particularly in Vietnam, Indonesia, and Congo. In the bar graph for K= 5 (Fig. 4), isolates from Brazil, Colombia and Uruguay clustered together (blue), China and the Congo cluster together (maroon), Indonesia (orange) and South Africa (green), each cluster independently and Malaysia, Oman, Pakistan and Vietnam cluster together (purple). Subsequent within-cluster STRUCTURE runs suggested K = 2 and 6 for cluster 1 (Supplementary Fig. 8b) and K = 2 and 9 for cluster 2 (Supplementary Fig. 8c). The observed substructures were congruent with the clusters observed in the analyses using all the data.

**Fig. 4.**
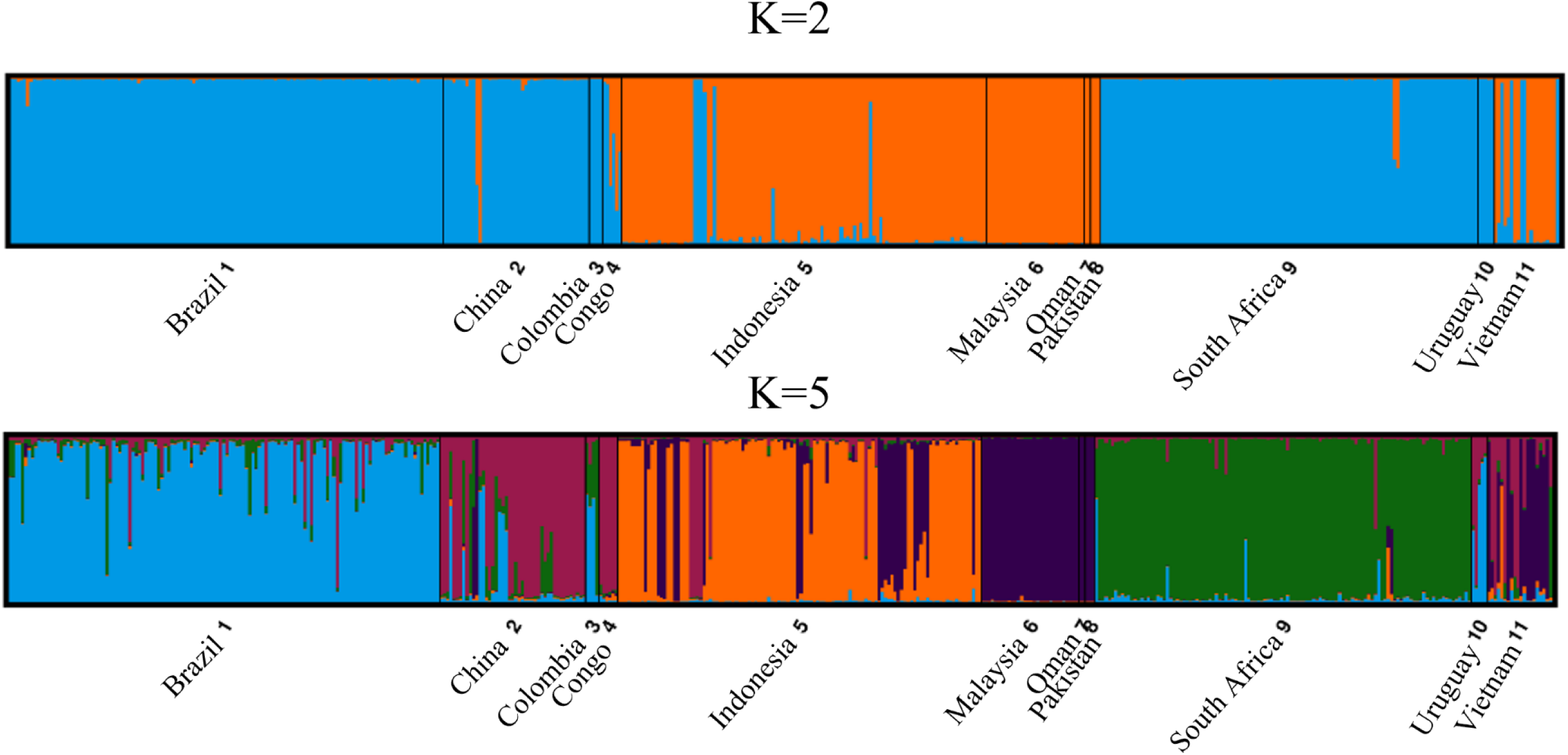
Analysis of population structure (K) of a clone-corrected dataset of the *Ceratocystis* isolates collected from11 countries. Each individual is represented by a single vertical line and the colours indicate the relatedness of an isolate to a specific cluster. The STRUCTURE analysis, with no prior assumptions, revealed an optimal number of 2 and 5 clusters.

The DAPC K-means analyses and the BIC clustering analysis on the non-clone-corrected dataset was congruent with the clusters identified by STRUCTURE analysis and separated the data into two genetic clusters (Fig. 5). Individuals from Brazil, Colombia, Congo, South Africa, and Uruguay were exclusive to cluster 1 and Malaysia, Oman, Pakistan were exclusive to cluster 2. Individuals from Vietnam and Indonesia predominately grouped with cluster 2 but individuals were found across both clusters. The inverse was true for China as individuals predominantly grouped with cluster 1 but were found across both clusters. The clear separation and tight within-cluster grouping indicate genetically distinct populations with limited genetic exchange or admixture between the two populations.

**Fig. 5.**
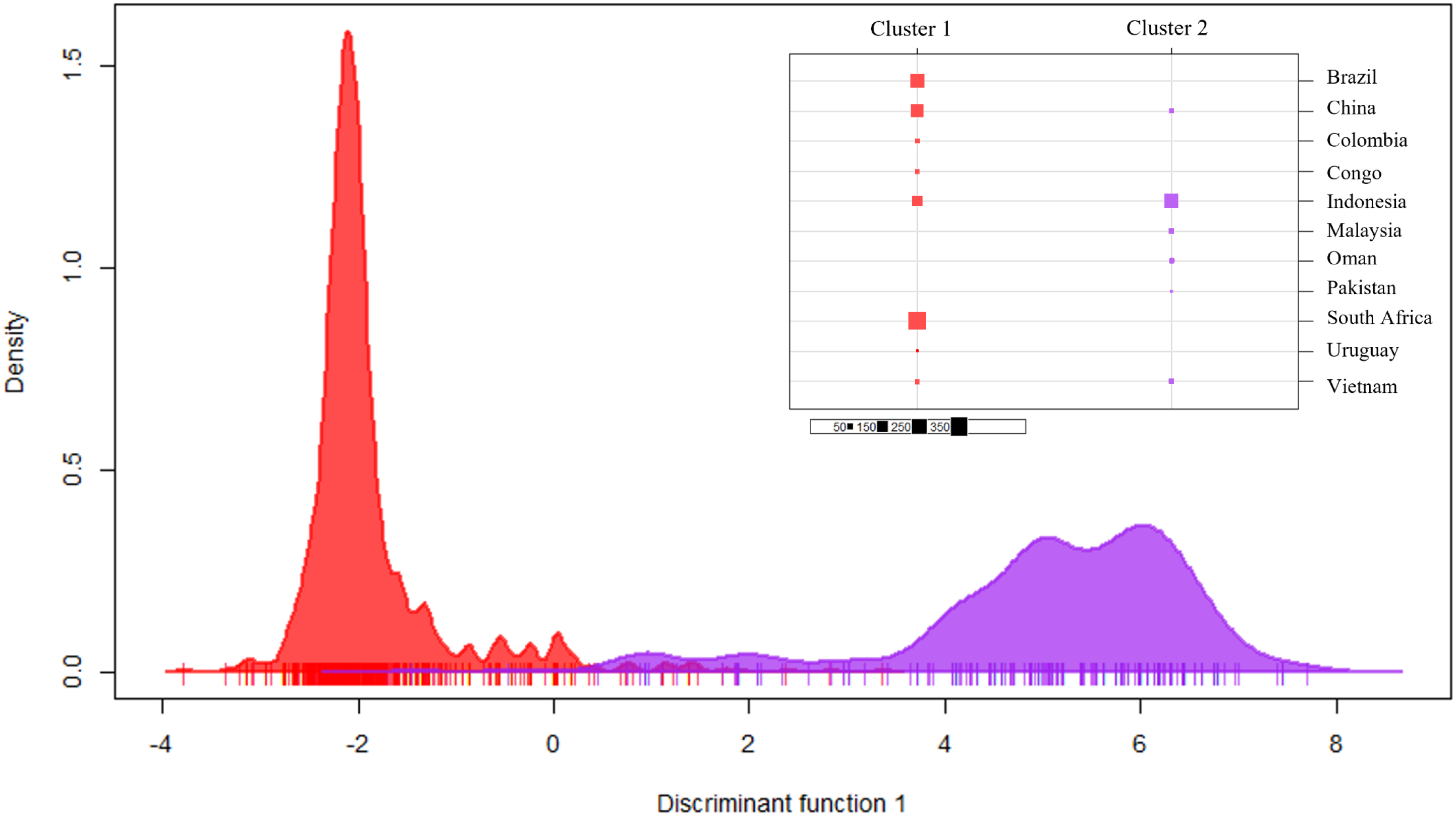
Discriminant Analysis of Principal Components (DAPC) of genetic clusters. DAPC plot showing the distribution of isolates in two distinct bell curves. Cluster 1 encompasses isolates from Brazil, Colombia, Congo, South Africa, and Uruguay, while Cluster 2 includes isolates from Malaysia, Oman, and Pakistan. Isolates from Vietnam and Indonesia predominantly group with Cluster 2 but are also present in both clusters. Conversely, isolates from China mainly group with Cluster 1 but are found across both clusters. The distinct bell curves and minimal overlap suggest that these populations are genetically distinct, with limited genetic exchange or admixture between them.

The DAPC K-means analyses, and the BIC clustering analysis on the isolates showing admixture resolved into two groups (Supplementary Fig. 9). Individuals from Indonesia were exclusive to group 1 and individuals from Brazil, Colombia and South Africa were exclusive to group 2.

### 3.6 Minimum spanning network analyses

The MSN analyses (Edwards distances) revealed two distinct clusters for the 1174 (Fig. 6) and were congruent with clusters found in ML, STRUCTURE and DAPC K-means analyses. Isolates from Brazil, Colombia, Congo, South Africa, and Uruguay were exclusive to cluster 1 (Lineage 1). These isolates were also more closely related to each other. Similarly, isolates from Malaysia, Oman, Pakistan were exclusively found in cluster 2 (Lineage 2). Isolates from Vietnam (Supplementary Fig. 10a) and Indonesia (Supplementary Fig. 10b) and China (Supplementary Fig. 10c) were found in both clusters, with isolates from Indonesia demonstrating greater differentiation between the haplotypes based on the thickness of connecting branches. Across the dataset, six MLGs were shared between countries. Oman and Pakistan shared a MLG, and Brazil and Colombia shared a MLG. China shared the most MLGs: one with Indonesia, one with South Africa, and two with Brazil. Of the shared MLGs, five were within cluster 1. Country-level MSNs are in Supplementary Fig. 10. The MSN for admixed isolates showed no structure nor shared haplotypes across populations. Admixed isolates from Indonesia and Colombia were more closley related to themselves and to each other, than to isolates from Brazil. Similarly isolates from South Africa were more closley related to those from Brazil Supplementary Fig. 11.

**Fig. 6.**
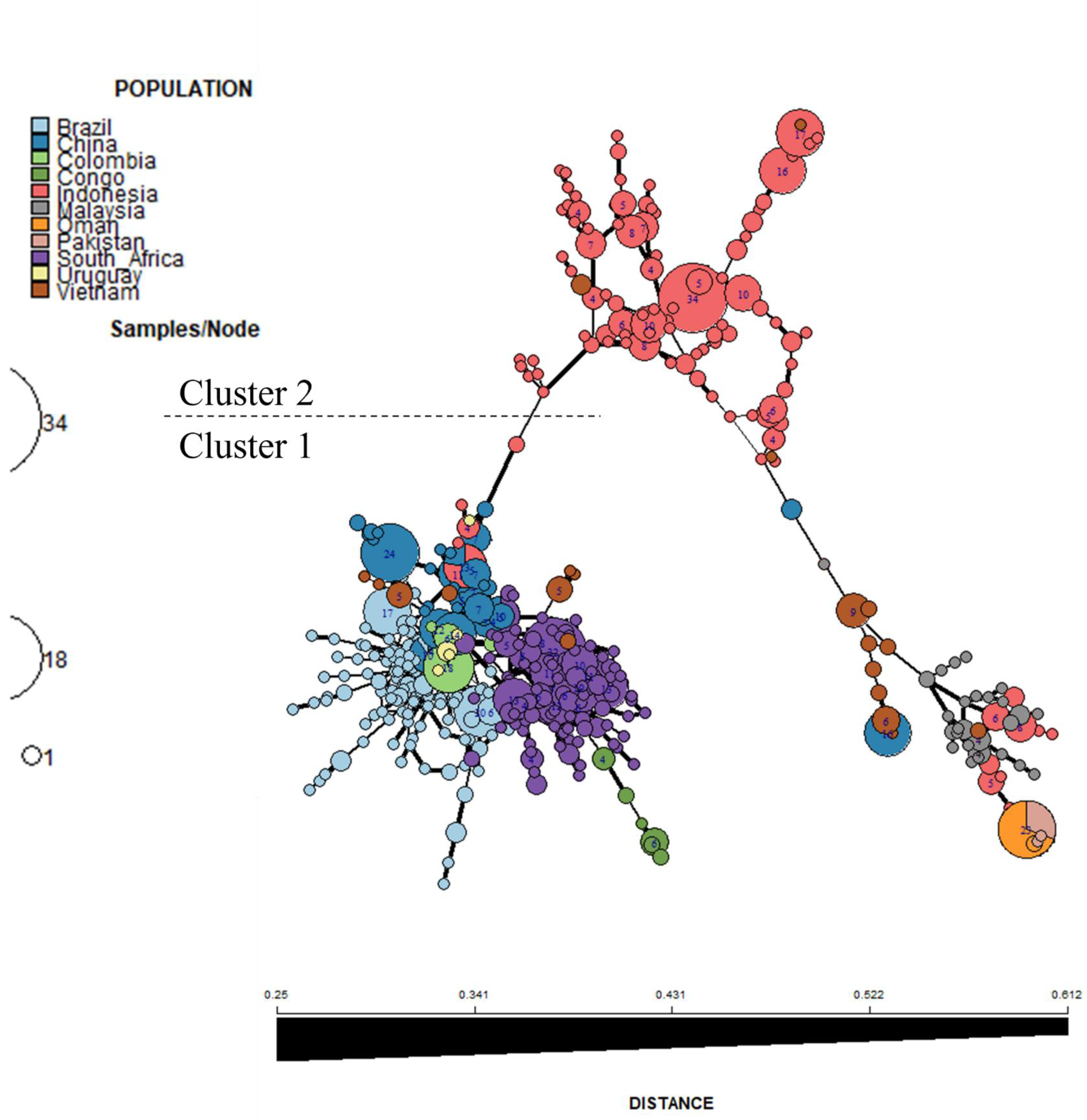
Minimum spanning network (MSN) showing the relationship of the *Ceratocystis* isolates based on Edwards genetic distance. Each node represents one multilocus genotype (MLG) and the size of the node is proportional to the number of individuals with that MLG. Nodes are coloured according to sampling location. The colour gradient of the lines between nodes indicates genetic distance: where dark lines denote closer genetic similarity and light lines denote greater divergence.

### 3.7 Pairwise population differentiation and gene flow

The clusters identified by STRUCTURE and DAPC were statistically supported by AMOVA calculations, which indicated significant between-population differentiation (44%) as well as 56% within-population variance (Table 2). These values indicate both high within-population diversity and strong among-population structure. AMOVA calculations for the isolates that indicated a mixed ancestry, indicated that there was significant differentiation between populations (64 %), where populations were defined by isolates per country.

**Table 2.**
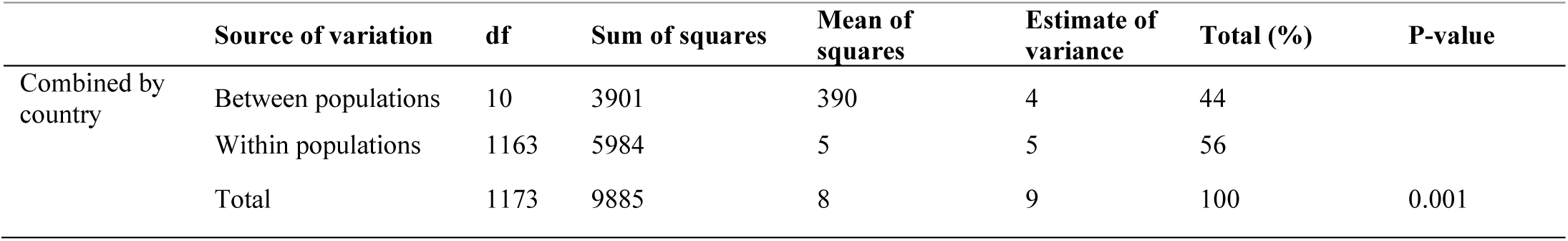
Hierarchical analysis of molecular variance (AMOVA) of *Ceratocystis* isolates grouped by populations based on country of origin.

Pairwise differentiation and gene flow analyses, using country-level populations (Table 3), showed lower Gst values when comparing countries whose isolates belonged to the same lineage, and higher values when comparing countries from different lineages. For example, Gst was near 1 (0.98) when comparing Brazil (Lineage 1) to Oman (Lineage 2), but lower (0.38) between Brazil and Colombia (both Lineage 1), indicating less differentiation. This pattern was mirrored in Nm values, with values close to zero reflecting high differentiation and limited gene flow. Results were consistent with ML, STRUCTURE, DAPC, and MSN analyses. For the two *Ceratocystis* sub-populations (Lineage 1: *C. eucalypticola*; Lineage 2: *C. manginecans*), Gst = 0.68 and Nm = 0.12 indicate high differentiation and limited gene flow, suggesting these lineages may be undergoing speciation driven by geographic isolation. All pairwise comparisons were significant (p = 0.000–0.021) based on 1,000 permutations.

**Table 3.**
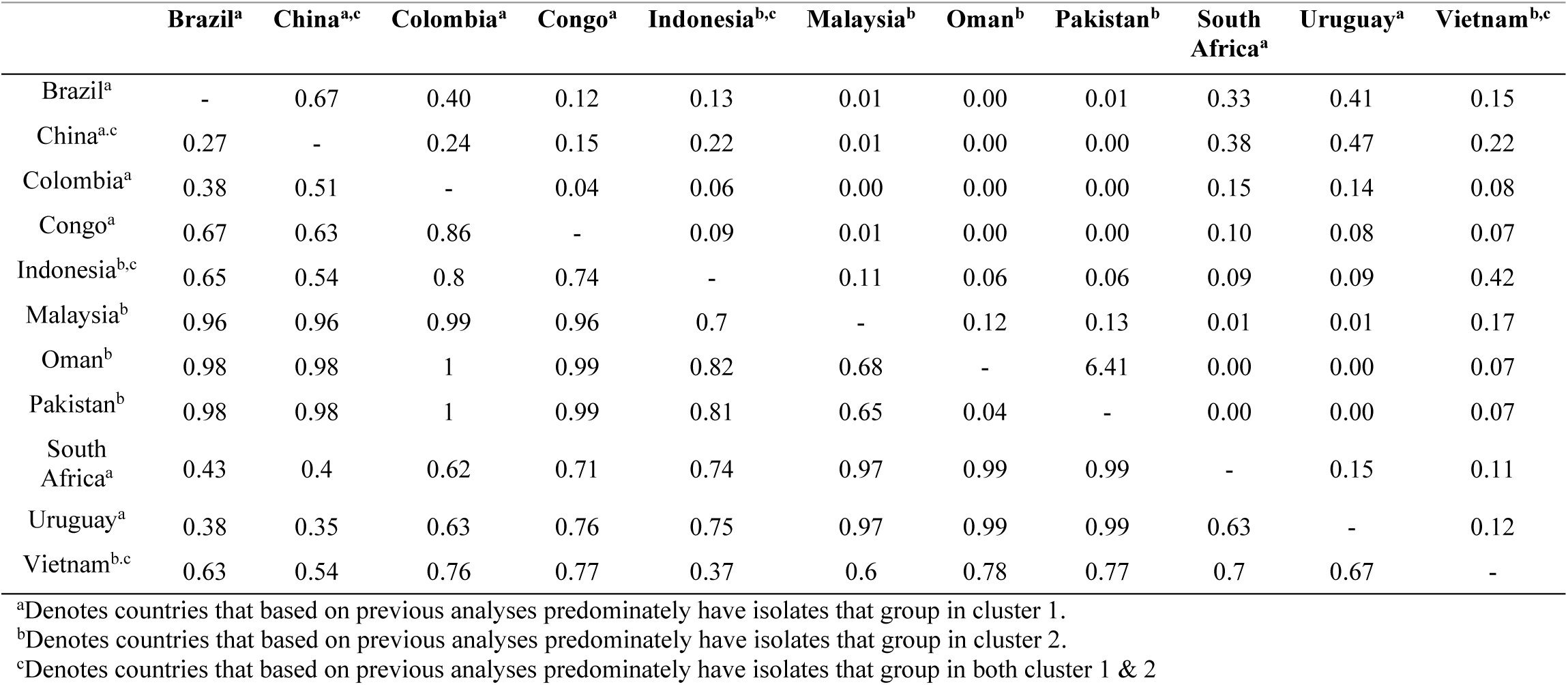
Population differentiation of all the *Ceratocystis* populations separated based on country. Pairwise comparison of population differentiation (ϕPT) using Hendricks standardized measure of genetic differentiation (Gst) and gene flow (Nm) among the populations are presented below and above the diagonal, respectively.

### 3.8 Morphological comparisons

The two-sample t-test rejected the null hypothesis in 19 of the 24 characters assessed, thus exhibiting significant differences between the two clusters (Table 4). For the statistically significant characters, the degree of difference was calculated as the difference of the averages of Lineage 1 and Lineage 2, divided by the average of both clusters, to provide a relative measure of the difference detected (absolute difference: Table 4). The average difference in dimensions between the two species ranged from 1 % to 12 % (Table 4). Notably, the ratio of branched to unbranched aleuriospore-conidiophores demonstrated a difference of 121 %, with Lineage 1 having a ratio of 0.19 compared to 0.77 in Lineage 2 (Fig. 7). Additionally, the average difference in the length of the ostiolar hyphae was 23 %, with measurements of 44.9 µm for Lineage 1 and 56.7 µm for Lineage 2.

**Table 4.**
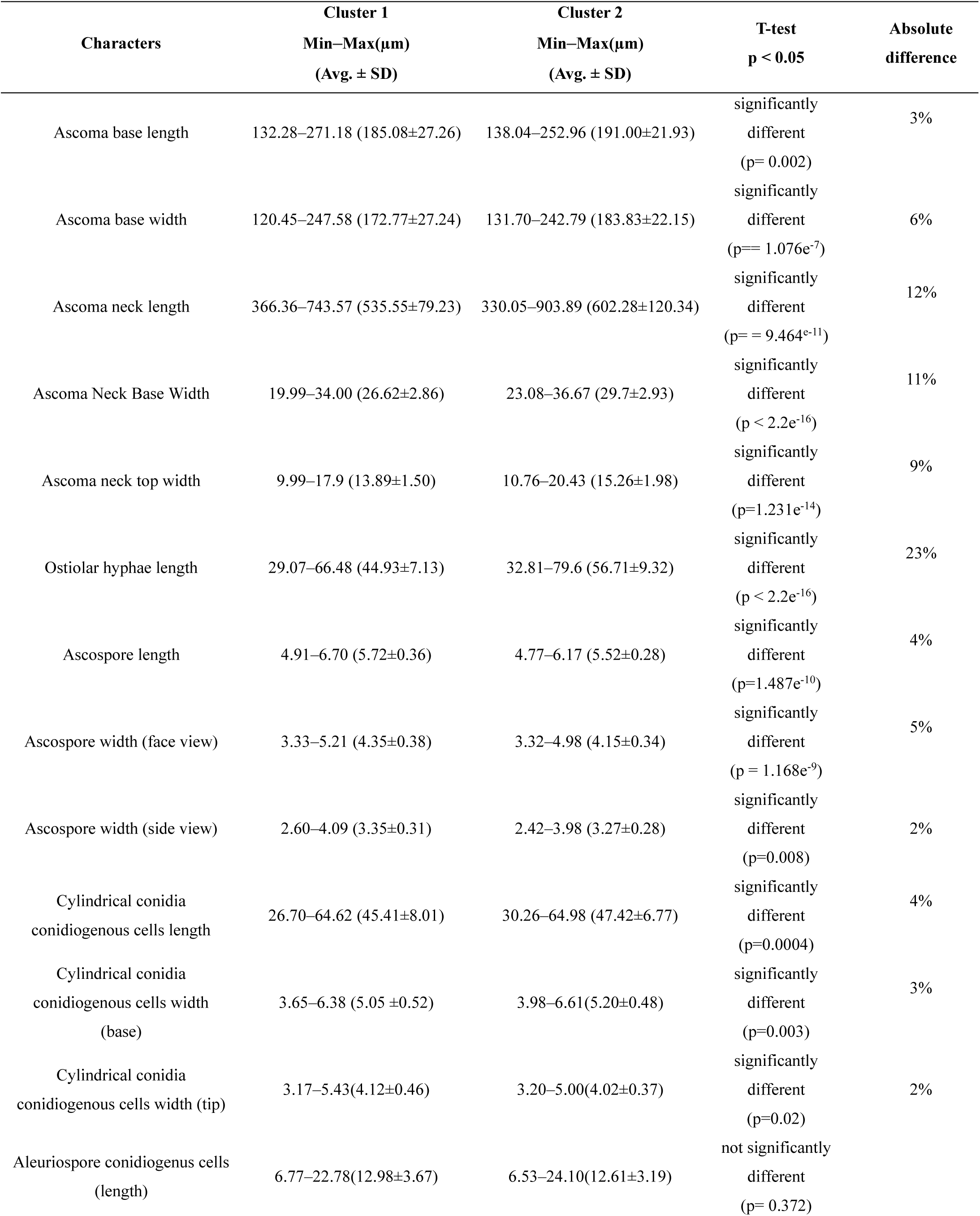
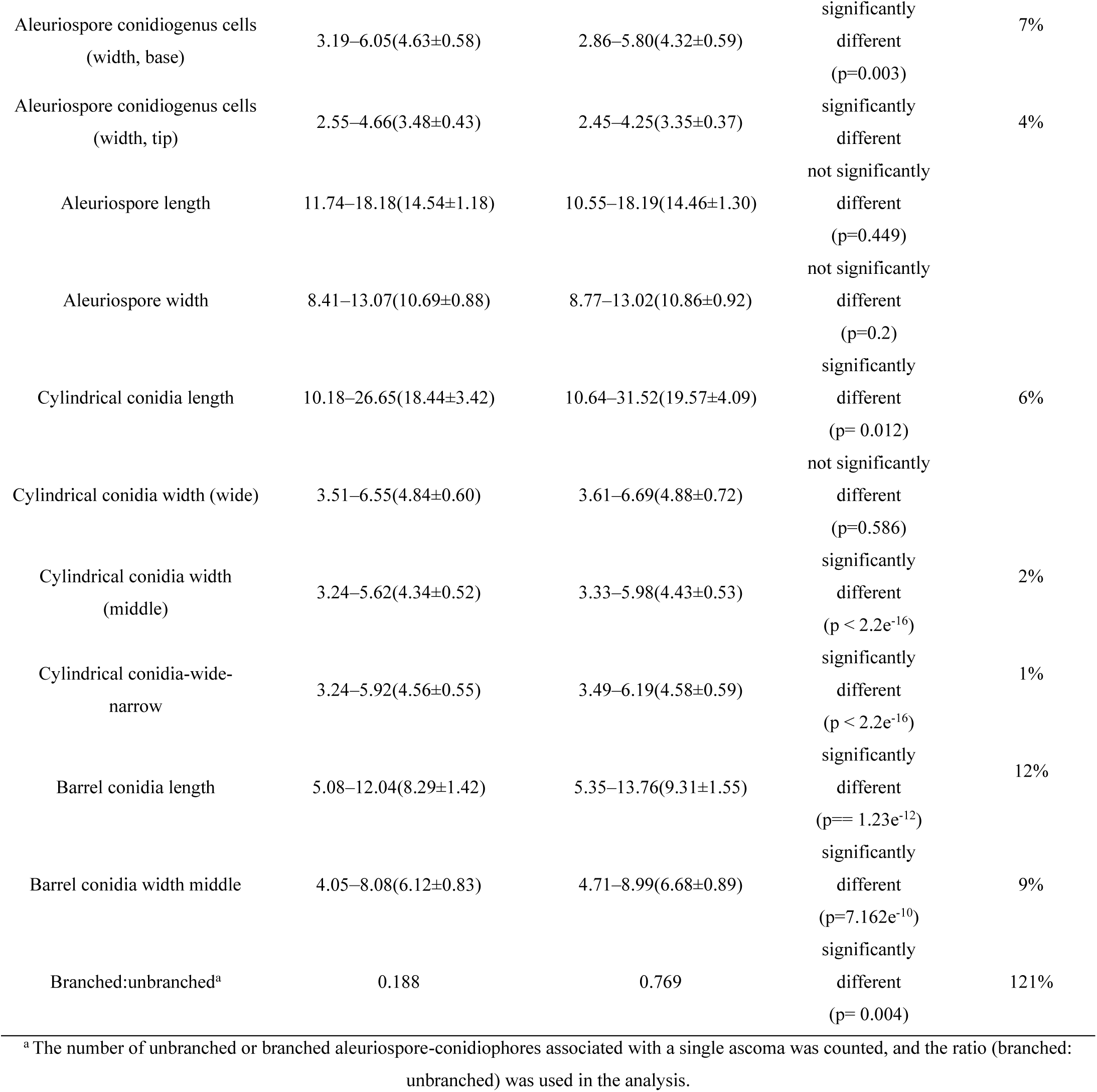
Comparative morphological characteristics, statistical analysis, and absolute differences quantifying the size of differences among 18 replicates of three *Ceratocystis* isolates per cluster: Cluster 1 for Lineage 1 and Cluster 2 for Lineage 2.

| Characters | Cluster 1<br>Min–Max(μm)<br>(Avg. ± SD) | Cluster 2<br>Min–Max(μm)<br>(Avg. ± SD) | T-test<br>p < 0.05 | Absolute<br>difference |
| --- | --- | --- | --- | --- |
| Ascoma base length | 132.28–271.18 (185.08±27.26) | 138.04–252.96 (191.00±21.93) | significantly<br>different<br>(p= 0.002) | 3% |
| Ascoma base width | 120.45–247.58 (172.77±27.24) | 131.70–242.79 (183.83±22.15) | significantly<br>different<br>(p= 1.076e<sup>-7</sup>) | 6% |
| Ascoma neck length | 366.36–743.57 (535.55±79.23) | 330.05–903.89 (602.28±120.34) | significantly<br>different<br>(p= 9.464e<sup>-11</sup>) | 12% |
| Ascoma Neck Base Width | 19.99–34.00 (26.62±2.86) | 23.08–36.67 (29.7±2.93) | significantly<br>different<br>(p < 2.2e<sup>-16</sup>) | 11% |
| Ascoma neck top width | 9.99–17.9 (13.89±1.50) | 10.76–20.43 (15.26±1.98) | significantly<br>different<br>(p=1.231e<sup>-14</sup>) | 9% |
| Ostiolar hyphae length | 29.07–66.48 (44.93±7.13) | 32.81–79.6 (56.71±9.32) | significantly<br>different<br>(p < 2.2e<sup>-16</sup>) | 23% |
| Ascospore length | 4.91–6.70 (5.72±0.36) | 4.77–6.17 (5.52±0.28) | significantly<br>different<br>(p=1.487e<sup>-10</sup>) | 4% |
| Ascospore width (face view) | 3.33–5.21 (4.35±0.38) | 3.32–4.98 (4.15±0.34) | significantly<br>different<br>(p = 1.168e<sup>-9</sup>) | 5% |
| Ascospore width (side view) | 2.60–4.09 (3.35±0.31) | 2.42–3.98 (3.27±0.28) | significantly<br>different<br>(p=0.008) | 2% |
| Cylindrical conidia<br>conidiogenous cells length | 26.70–64.62 (45.41±8.01) | 30.26–64.98 (47.42±6.77) | significantly<br>different<br>(p=0.0004) | 4% |
| Cylindrical conidia<br>conidiogenous cells width<br>(base) | 3.65–6.38 (5.05 ±0.52) | 3.98–6.61(5.20±0.48) | significantly<br>different<br>(p=0.003) | 3% |
| Cylindrical conidia<br>conidiogenous cells width (tip) | 3.17–5.43(4.12±0.46) | 3.20–5.00(4.02±0.37) | significantly<br>different<br>(p=0.02) | 2% |
| Aleuriospore conidiogenous cells<br>(length) | 6.77–22.78(12.98±3.67) | 6.53–24.10(12.61±3.19) | not significantly<br>different<br>(p= 0.372) |  |
| Aleuriospore conidiogenous cells<br>(width, base) | 3.19–6.05(4.63±0.58) | 2.86–5.80(4.32±0.59) | significantly<br>different<br>(p=0.003) | 7% |
| Aleuriospore conidiogenous cells<br>(width, tip) | 2.55–4.66(3.48±0.43) | 2.45–4.25(3.35±0.37) | significantly<br>different | 4% |
| Aleuriospore length | 11.74–18.18(14.54±1.18) | 10.55–18.19(14.46±1.30) | not significantly<br>different<br>(p=0.449) |  |
| Aleuriospore width | 8.41–13.07(10.69±0.88) | 8.77–13.02(10.86±0.92) | not significantly<br>different<br>(p=0.2) |  |
| Cylindrical conidia length | 10.18–26.65(18.44±3.42) | 10.64–31.52(19.57±4.09) | significantly<br>different<br>(p = 0.012) | 6% |
| Cylindrical conidia width (wide) | 3.51–6.55(4.84±0.60) | 3.61–6.69(4.88±0.72) | not significantly<br>different<br>(p=0.586) |  |
| Cylindrical conidia width<br>(middle) | 3.24–5.62(4.34±0.52) | 3.33–5.98(4.43±0.53) | significantly<br>different<br>(p < 2.2e<sup>-16</sup>) | 2% |
| Cylindrical conidia-wide-<br>narrow | 3.24–5.92(4.56±0.55) | 3.49–6.19(4.58±0.59) | significantly<br>different<br>(p < 2.2e<sup>-16</sup>) | 1% |
| Barrel conidia length | 5.08–12.04(8.29±1.42) | 5.35–13.76(9.31±1.55) | significantly<br>different<br>(p== 1.23e<sup>-12</sup>) | 12% |
| Barrel conidia width middle | 4.05–8.08(6.12±0.83) | 4.71–8.99(6.68±0.89) | significantly<br>different<br>(p=7.162e<sup>-10</sup>) | 9% |
| Branched:unbranched<sup>a</sup> | 0.188 | 0.769 | significantly<br>different<br>(p= 0.004) | 121% |

<sup>a</sup> The number of unbranched or branched aleuriospore-conidiophores associated with a single ascoma was counted, and the ratio (branched: unbranched) was used in the analysis.

**Fig. 7.**
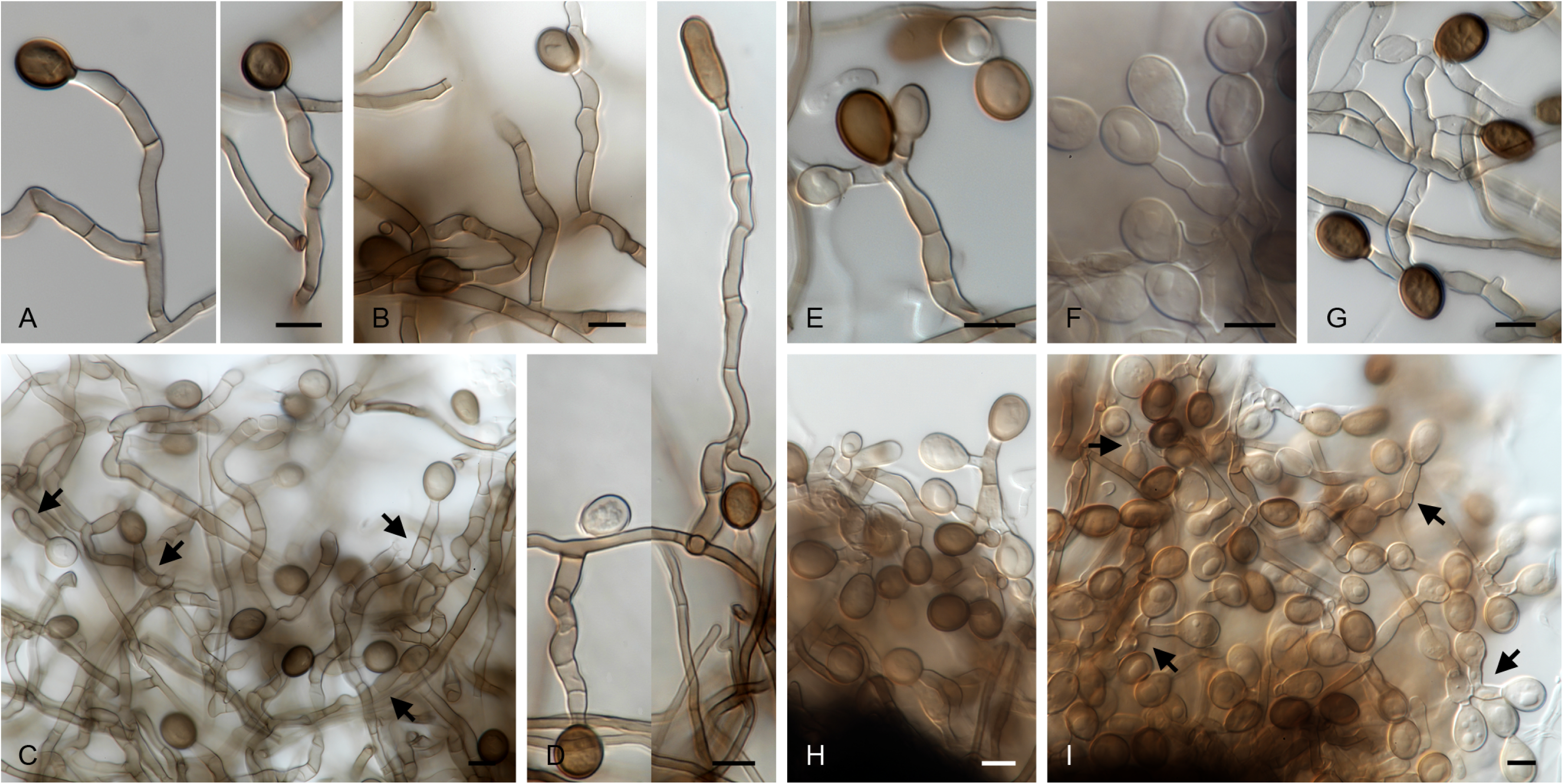
Branching pattern of conidiophores of Lineage 1 (CMW:56740 (A, C), CMW:57998 (B)). and Lineage 2 (CMW:60227 (E–I), CMW:60230 (D)). A–D: unbranched conidiophores. E–I: branched conidiophores. Scale bars A–I = 10 µm.

## 4. Discussion

This study considered a large number of *Ceratocystis* isolates from historical and new collections across 11 countries and eight host species including *Acacia*, *Eucalyptus* and *Mangifera*. Isolates from historical outbreaks of Ceratocystis wilt and canker had previously been identified as *C. manginecans*, *C. eucalypticola*, or residing in the *C. fimbriata sensu lato* or *C. manginecans* complexes. Using this extensive dataset, phylogenetic analyses were conducted for seven gene regions and population genetic assessments were made using 16 microsatellite markers to consider possible species boundaries. All analyses consistently grouped the data for historical and newly acquired isolates into two distinct lineages within the *C. manginecans* complex as defined by Harrington et al., (2023). One of these represented isolates previously designated as *C. eucalypticola* and the other those previously designated as *C. manginecans*. Morphological comparisons between representatives of each of the two lineages showed some distinct characteristics. The study also revealed limited gene flow between isolates defining the two lineages but also showed evidence of hybridization between them.

Multi-gene phylogenetic analyses supported the delineation of the newly acquired isolates into two statistically supported lineages within the *C. manginecans* complex. Phylogenetic analyses of sequence data for four gene regions revealed that one set of isolates formed a monophyletic clade, identified in this study as Lineage 1 including the isolate previously designated as the ex-type of *C. eucalypticola.* A second set of isolates formed a monophyletic clade that we refer to as Lineage 2, that included the ex-type isolate of *C. manginecans*. The strongest signals distinguishing these two lineages were from sequences of the coding genes rpb2 and MS204, which, in combination, differentiate 13 lineages in the LAC. Sequence data for the MAT gene regions were included in this study to define species boundaries in *Ceratocystis*, as recommended by Harrington et al., (2023). Those results showed that the clustering of isolates using ML analyses of the MAT region was largely congruent with the multi-gene ML phylogenies. This was apart from some isolates, that clustered inconsistently across the two MAT ML phylogenies. Thus, while sequences for the MAT region complement multi-gene analyses, on their own, they are insufficient to fully resolve species boundaries.

Significant variability was identified in the ITS region among the screened isolates, with those from Brazil exhibiting the highest number of distinct ITS sequence variants (haplotypes). This intraspecific and intragenomic variation in rDNA indels is well recognised and renders ITS sequences unreliable for species delineation (Harrington et al., 2014; Naidoo et al., 2013). The results highlight the complexities of defining species boundaries based solely on representative genotypes sampled from a population and underscore the importance of understanding natural variation within and among populations for accurate species delineation (Oliveira et al., 2015).

Population genetic analyses for 1174 isolates utilising 16 microsatellite markers, revealed two distinct population clusters with limited gene flow between them. These were consistent with the two lineages emerging from the phylogenetic analyses for isolates previously designated as *C. eucalypticola* (Lineage 1) and *C. manginecans* (Lineage 2). Similar to the results of Liu et al. (2021), isolates in population cluster 1 were predominantly from *Eucalyptus* and had a greater geographic distribution but were less genetically diverse. In contrast, isolates in population cluster 2 were confined to Asia but exhibited greater genetic diversity and were collected from a wider variety of hosts. The population from Vietnam was the most genetically diverse, which is consistent with the results of Fourie et al., (2016). However, at a sub-population level, analyses for population cluster 1 showed that isolates from Brazil were most genetically diverse, which is consistent with results of Ferreira et al., (2011). Limited gene flow was evident between the population clusters, suggesting that while they represent distinct entities, they are not fully reproductively isolated. This implies that these lineages may have different native ranges or that they may have been introduced to these regions long ago and are now undergoing speciation driven by host associations and geographic isolation (Harrington et al., 2023; Liu et al., 2021).

Both phylogenetic and population genetic analyses provided evidence of hybridization between isolates in the two lineages identified in this study and previously identified as *C. eucalypticola* and *C. manginecans* in several regions. Contrasting gene topologies indicated mixed ancestry resulting from hybridization events, a phenomenon previously reported in *Ceratocystis* spp. (Engelbrecht and Harrington, 2005; Kanzi et al., 2020; Naidoo et al., 2013) and other fungal species complexes (Sillo et al., 2019). In this study, isolates with mixed ancestry were identified in Brazil, Colombia, South Africa, and Indonesia, primarily from commercially propagated *Eucalyptus* clones. While no shared haplotypes were found between populations, DAPC analysis revealed two clusters. One of these included hybrid isolates from Brazil, Colombia, and South Africa, exhibiting contrasting gene topologies in the ITS and MS204 regions, suggesting an ancestral hybridization event followed by a likely dispersal through trade in *Eucalyptus* products (Ferreira et al., 2013, 2011; Liu et al., 2021). The second cluster consisted of isolates from Indonesia, collected from *Eucalyptus* clones and an *Acacia* sp., indicating a potential separate hybridization event, possibly due to the presence of isolates from the two different lineages in the region. Several of these individuals exhibited contrasting gene topologies in the rpb2 and MS204 regions. These findings suggest limited but ongoing gene flow between these recently diverged lineages, consistent with the idea that gene flow will continue until reproductive barriers become established (Harrison and Larson, 2014).

Results of this study provided some insights into the pathways of spread of isolates residing in the two identified lineages in the *C. manginecans* complex. Regions with lower genetic diversity, such as Colombia, suggest a recent introduction of Lineage 1 (population cluster 1). This introduction has likely been via infected *Eucalyptus* propagation material, supported by the shared cluster 1 MLG (collected from *Eucalyptus* clones) between Colombia and Brazil. The spread of isolates in Lineage 1 (population cluster 1) via the exchange of *Eucalyptus* propagation material has previously been suggested as a pathway of introduction (Ferreira et al., 2013, 2011; Liu et al., 2021). It is also supported in the results of this study, with the majority of shared MLGs identified in this study collected from *Eucalyptus* clones and residing in population cluster 1 (Lineage 1). Another important pathway of spread, particularly for Lineage 2 (population cluster 2), appears to be through host expansion events. This has been demonstrated by genetic associations between *Ceratocystis* (cluster 1) on *Punica* and *Eucalyptus* (Harrington et al., 2014; Liu et al., 2021) and *Acacia* and *Eucalyptus* in Indonesia (cluster 2: Hlongwane, 2022). A similar host expansion of lineage 2 (population cluster 2) from *Acacia* to *Eucalyptus* was identified in Malaysia in this study. These findings highlight the critical pathways for pathogen spread and underscore the need for stringent quarantine measures.

The results of this study identified *Ceratocystis* isolates from previously unreported regions and hosts. Newly acquired isolates from Colombia were identified as residing in the *C. manginecans* complex grouping with lineage 1 (previously designated as *C. eucalypticola*), which is the first report of the lineage from *Eucalyptus* sp. in that country. These isolates were collected from freshly made wounds on *Eucalyptus* in the absence of any disease symptoms, similar to a situation reported form China (Liu et al., 2021) and South Africa (van Wyk et al., 2012). Furthermore, single isolates from *Mimusops elengi* and *Lansium domesticum* also resided in *C. manginecans* complex grouping with lineage 1 (previously designated as *C. eucalypticola*). Previously, these isolates had been identified as *C. manginecans* (Pratama et al., 2021b) and *C. fimbriata* (Suwandi et al., 2021), respectively. However, based on the results of the present study utilizing multiple gene regions, they should now be treated as Lineage 1 in the *C. manginecans* complex. Furthermore, isolates from *Eucalyptus* in Malaysia were identified as part of the *C. manginecans* complex grouping with Lineage 2 (previously designated as *C. manginecans*), which is the first report of this pathogen affecting *Eucalyptus* spp. in that country.

Field observations suggest potential differences in pathogenicity between the two lineages resolved in this study. Lineage 1 (formerly *C. eucalypticola*) has been recovered from fresh wounds on asymptomatic *Eucalyptus* in China (Liu et al., 2021), South Africa (van Wyk et al., 2012) and in Colombia (current study), suggesting that isolates can be only weakly pathogenic or even saprobic. In contrast, Lineage 2 (*C. manginecans*) has been reported only as the causal agent of aggressive wilt and canker disease on variety of different plant hosts (Al Adawi et al., 2013; Tarigan et al., 2011). Similar intra-specific variation in disease severity has been reported for other *Ceratocystis* pathosystems (Oliveira et al., 2016, 2021), underscoring the need for targeted inoculation trials to confirm and quantify the aggressiveness of these isolates. Such experiments, using representative isolates from both lineages and multiple host genotypes, are needed to test this hypothesis and guide breeding and quarantine strategies.

Geographic segregation and varying genetic diversities have important implications for quarantine and management strategies. Results of this study suggest that Lineage 1 (population cluster 1) and Lineage 2 (population cluster 2) have distinct geographical distributions. This has important quarantine consequences. Preventing the introduction of these pathogens into new regions will be an important first step in their management, reducing the likelihood of bridgehead effects (Lombaert et al., 2010), host expansion (Slippers et al., 2005), and hybridisation events (Brasier, 2000). If these pathogens successfully establish in new regions, monitoring the genetic diversity of these outbreaks becomes a crucial next step for managing them, as it informs resistant breeding strategies as demonstrated by Fernandes et al. (2024) and Oliveira et al. (2021).

A detailed comparison of isolates representing the two lineages identified in the *C. manginecans* complex, and from root or above-ground infections, respectfully revealed morphological distinctions between them. Although the morphological differences were subtle, the large sample size considered, showed that the differences were statistically significant in 19 of 24 characters analysed. The most notable of these differences was in the ratio of branched aleuriospore-conidiophores with isolates from Lineage 2 exhibiting significantly higher ratios. While morphological differences typically provide limited taxonomic resolution when considering closely related species (de Beer et al., 2014) they can add value. This was true in the current study where such differences aligned closely with phylogenetic and population genetic analyses. The ecological implications of these morphological variations merit further consideration, particularly in understanding how they may reflect distinct disease biology associated with above-ground and below-ground infections.

This study provided robust evidence for diverging lineages within the *C. manginecans* complex. There has been a long history of disagreement regarding the taxonomic position of these isolates, mostly arising from different species concepts applied to members of the LAC of *Ceratocystis*. However, phylogenetic analyses across seven gene regions, showed strong statistical evidence that isolates screened in this study, group into two distinct linages within the within the *C. manginecans* complex. Combined with genotype data for numerous loci and a large set of isolates, compelling evidence emerged supporting the divergence of lineages in the *C. manginecans* complex. Although they are not fully reproductively isolated distinguishing between them should facilitate communication within the research community and improve quarantine and disease management strategies.

This study adopted a taxonomic framework proposed by Harrington et al. (2023) in how the isolates were analysed. Following this approach *C. fimbriata* is recognised as a clonal lineage restricted to sweet potato (Harrington et al. 2023; Marincowitz et al., 2020), and the interfertile taxa *C. manginecans*, *C. eucalypticola*, *C. mangivora* and *C. mangicola* are placed within the *C. manginecans* complex (Harrington et al. 2023). Isolates in this complex are also interfertile with *C. fimbriata* and for this reason, some authors have treated them as *C. fimbriata sensu lato* (Fernandes et al., 2024). Although both viewpoints have merit, the concurrent use of both names could be confusing, and, for the present, a consensus to following the approach suggested by Harrington et al. (2023) would be least disruptive. However, further multilocus studies are required for other previously considered taxa recently placed in the *C. manginecans* complex, including *C. curvata*, to clarify their placement and relationships. The multilocus phylogenies in this study clearly separate the *C. fimbriata* clonal lineage from the two lineages representing isolates in the *C. manginecans* complex (Lineage 1 and Lineage 2), which also display differences in host range, geographic distribution and biology. Recognising these two lineages within the *C. manginecans* complex should provide a practical basis for diagnostics, epidemiology and quarantine while acknowledging their shared ancestry and partial reproductive compatibility.

## Data availability statement

All data are openly available in the supplementary materials. Sequence data have been deposited in the public repository NCBI, with accession numbers provided in the supplementary table 1.

## Acknowledgements

We thank Johannes Joubert for aiding with the statistical analyses in the morphological comparisons. We are grateful to Dr. Carlos Rodas for supplying isolates from Colombia and collaborators at the Department of Plant Pathology, Universidade Federal de Viçosa, Minas Gerais, Brazil, for providing the DNA samples for Brazilian isolates. Dr Marthin Tarigan is thanked for supplying the two isolates from *Lansium domesticum* and *Mimusops elengi.* This study was initiated through the bilateral agreement between the Forestry and Agricultural Biotechnology Institute (FABI), University of Pretoria, and the APRIL Group, RGE, Indonesia. We acknowledge funding from the RGE-FABI Tree Health Programme and the National Research Foundation (NRF, MND190619448979), South Africa.

## Author Contributions

Kira M.T. Lynn: Conceptualization, Investigation, Methodology, Formal analysis, Validation, Writing - original draft. Michael J. Wingfield: Conceptualization, Methodology, Resources, Funding, Project administration, Writing - review & editing, Supervision. Leonardo S. S. Oliveira: Resources, Writing - review & editing. Acelino C. Alfenas: Resources, Writing - review & editing. Rafael Ferreira Alfenas: Resources, Writing - review & editing. Seonju Marincowitz: Investigation, Methodology, Formal analysis, Validation, Writing - review & editing. Irene Barnes: Conceptualization, Methodology, Funding, Project administration, Writing - review & editing, Primary supervision.

## Supplementary data

**Supplementary Table 1.** Metadata for the *Ceratocystis* isolates investigated in this study. The table provides detailed information on the collection locations, the individuals who processed the samples, and includes all SSR and phylogenetic data for the gene regions analysed. File too large to display please find data in: Please see separate excel spread sheet.

**Supplementary Table 2.**
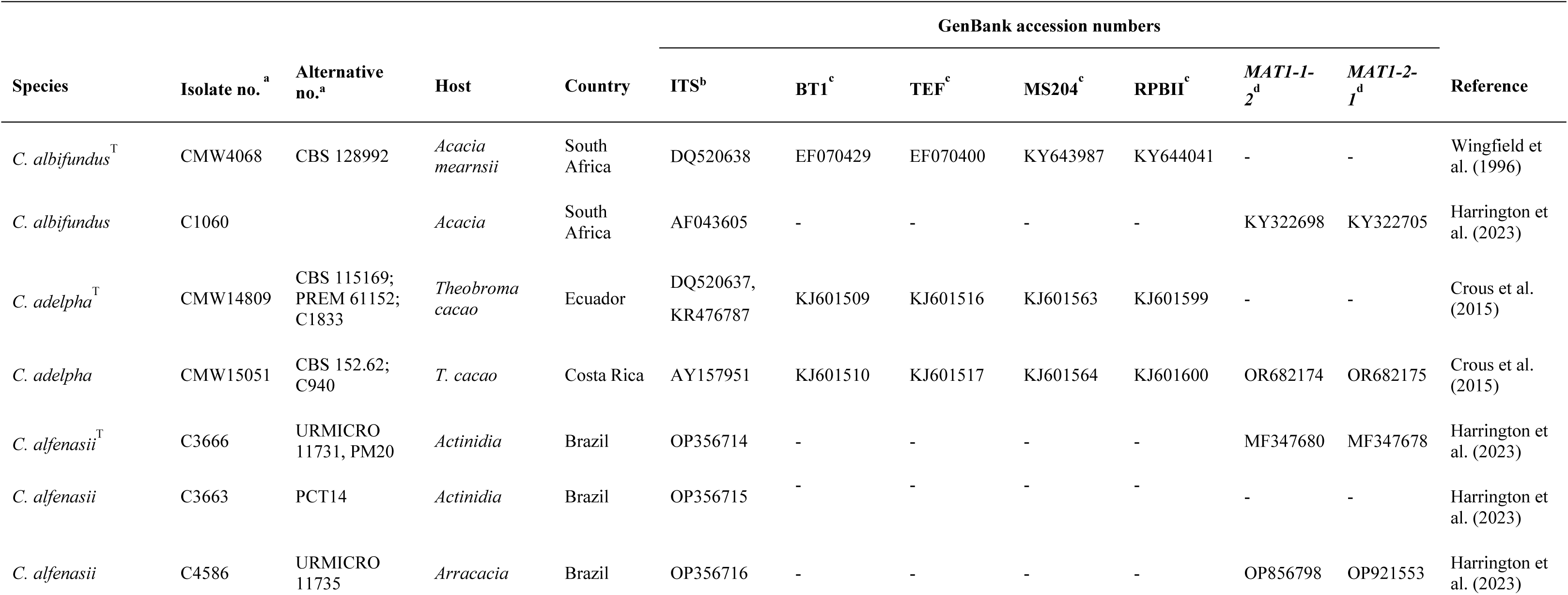
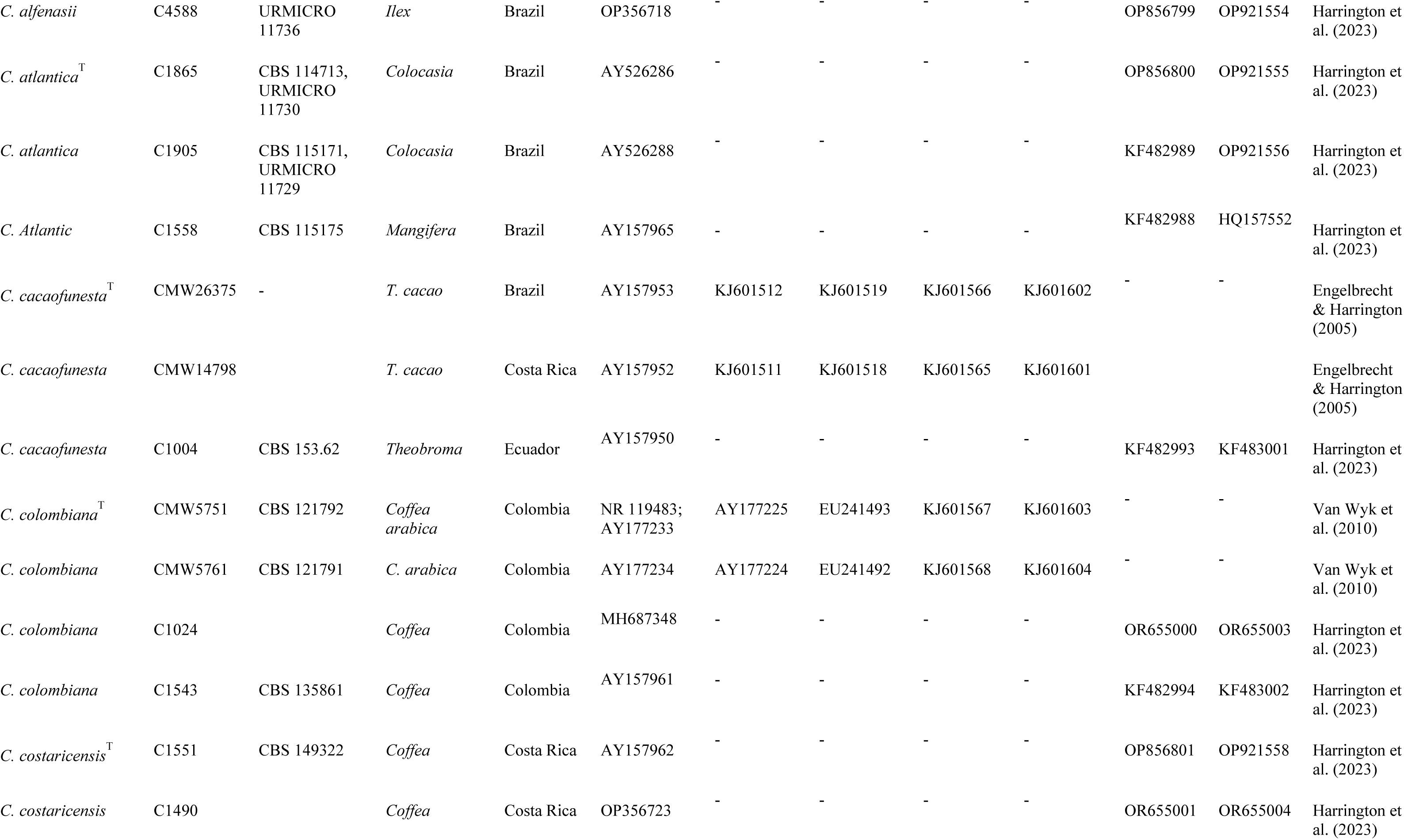
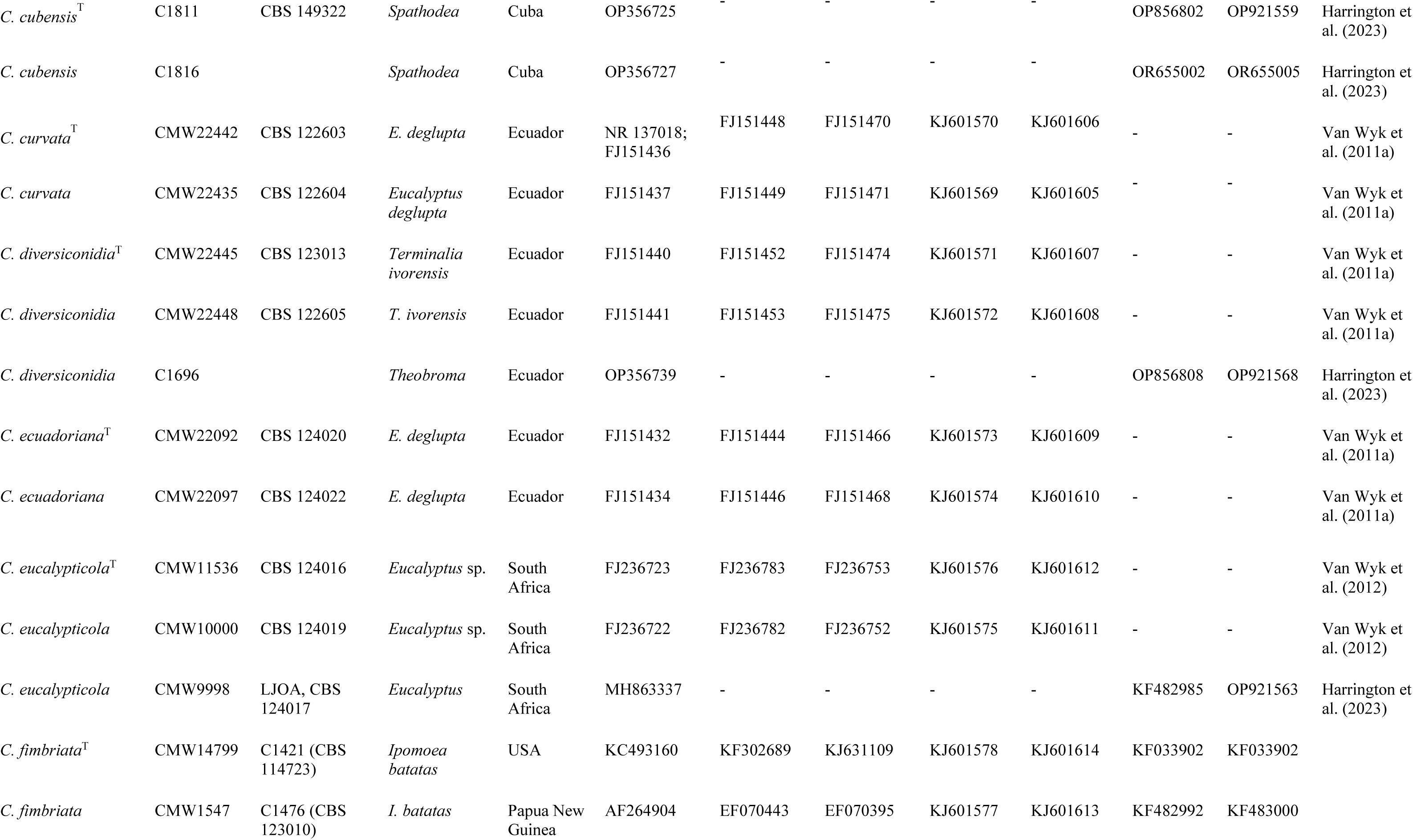
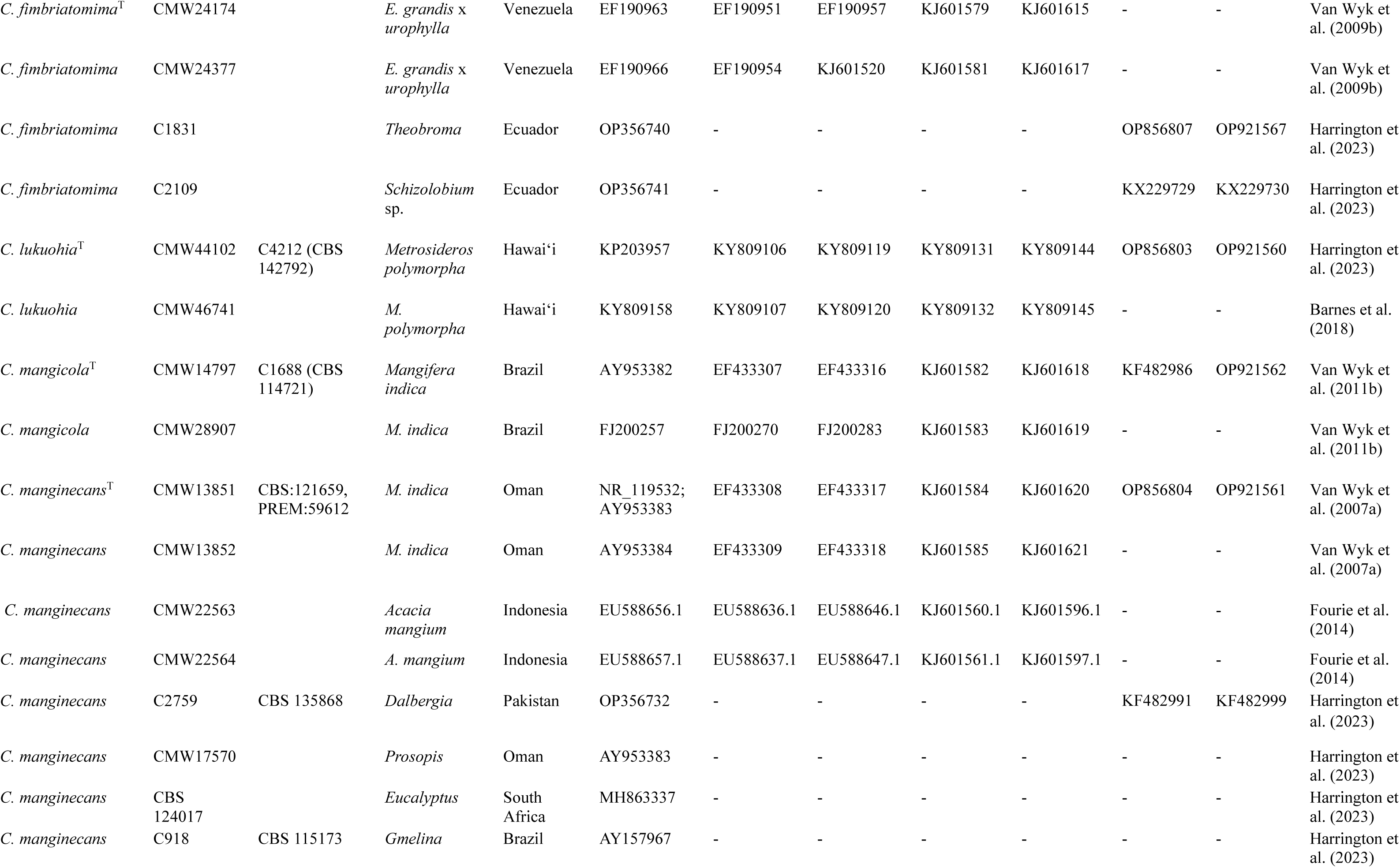
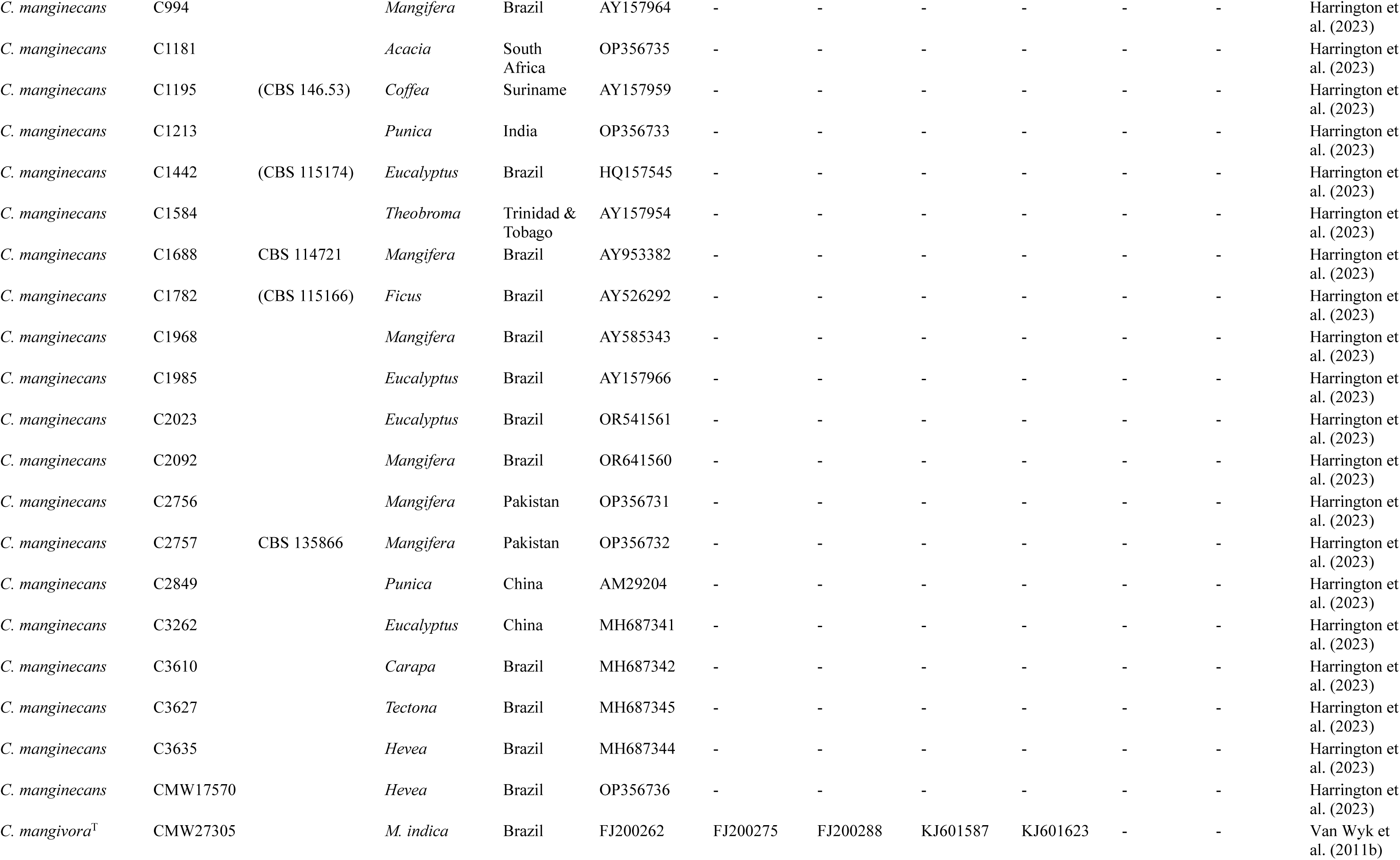
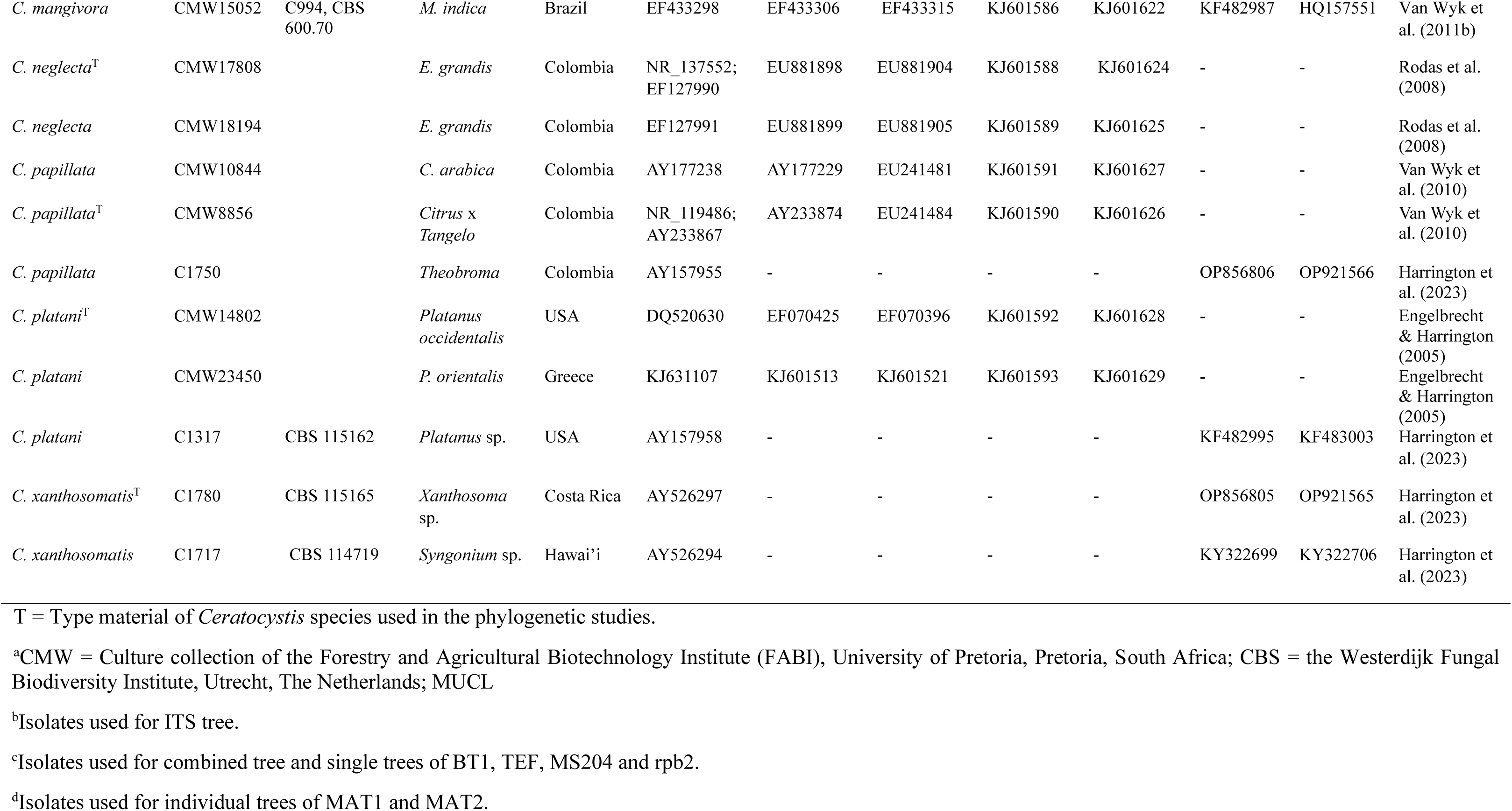
Details of the *Ceratocystis* isolates used for phylogenetic analysis in this study.

**Supplementary Fig. 1.**
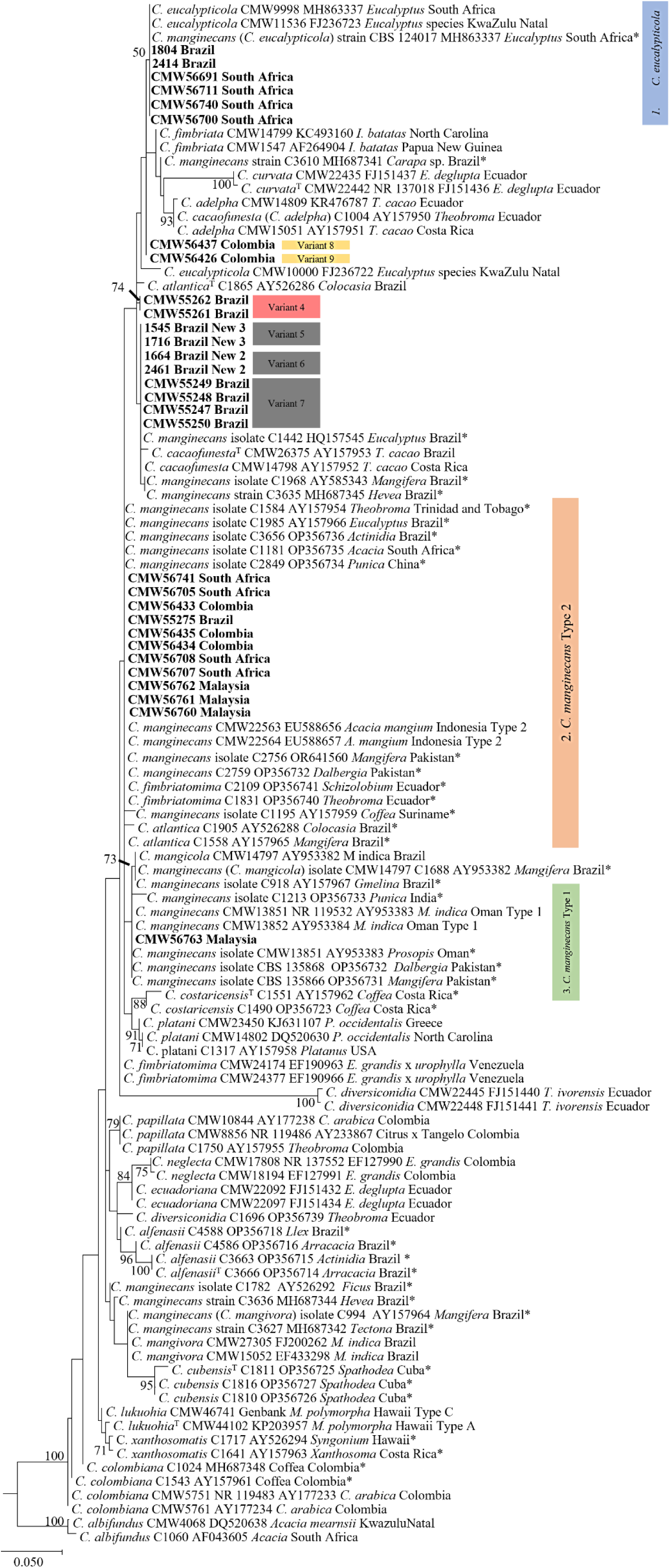
Phylogenetic tree based on maximum likelihood (ML) analysis of ITS sequences for *Ceratocystis* species in the Latin American Clade (LAC) and the *Ceratocystis* isolates used in this study (only representative haplotypes per country are shown in bold). Coloured boxes highlight the nine ITS sequence variants identified in this study from five regions: Brazil, Colombia, South Africa, Malaysia, and Indonesia. Although several variants (Variants 5–7, indicated in black, and Variants 8 & 9, indicated in yellow) cluster together with low statistical support, fixed SNP variations were observed in multiple isolates, forming distinct ITS variants. Bootstrap values above 50% are shown. For details regarding specific isolates please refer to the Supplementary Table 1. * Indicate isolates designated by Harrington et al. (2023).

**Supplementary Fig. 2.**
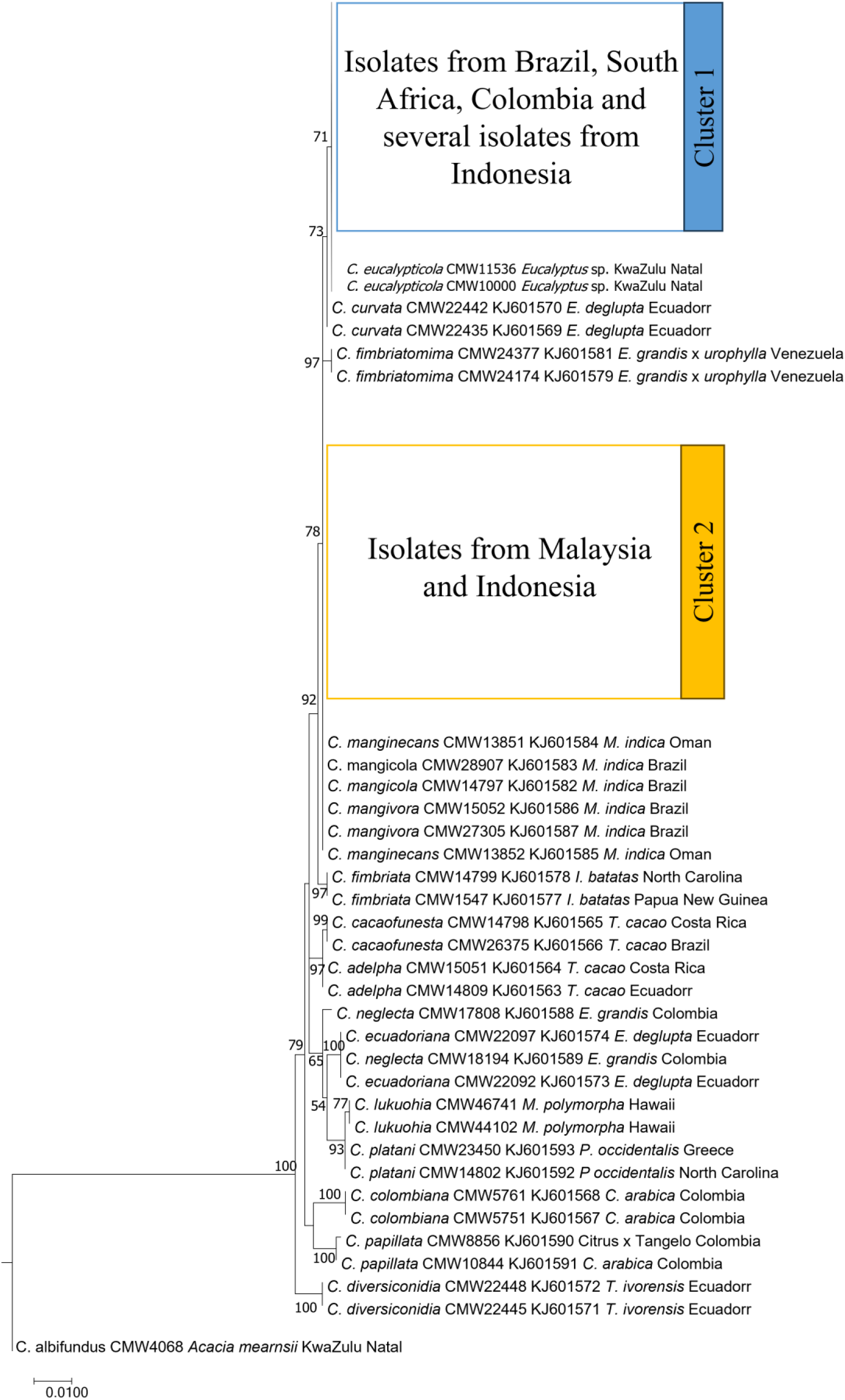
Phylogenetic tree based on maximum likelihood (ML) analysis of MS204 sequences for *Ceratocystis* species in the LAC and *Ceratocystis* isolates used in this study. Coloured boxes indicate representatives of isolates sequenced in this study from five regions (Brazil, Colombia, South Africa, Malaysia, and Indonesia). Isolates from Brazil, South Africa, Colombia and several isolates from Indonesia screened in this study formed a statistically supported monophyletic clade with Lineage 1 (*C. eucalypticola*: blue). Isolates from Malaysia and Indonesia clustered with Lineage 2 (*C. manginecans*: yellow) and isolates previously designated as *C. mangicola* and *C. mangivora*. Bootstrap values above 50% are shown. For details regarding specific isolates please refer to the Supplementary Table 1.

**Supplementary Fig. 3.**
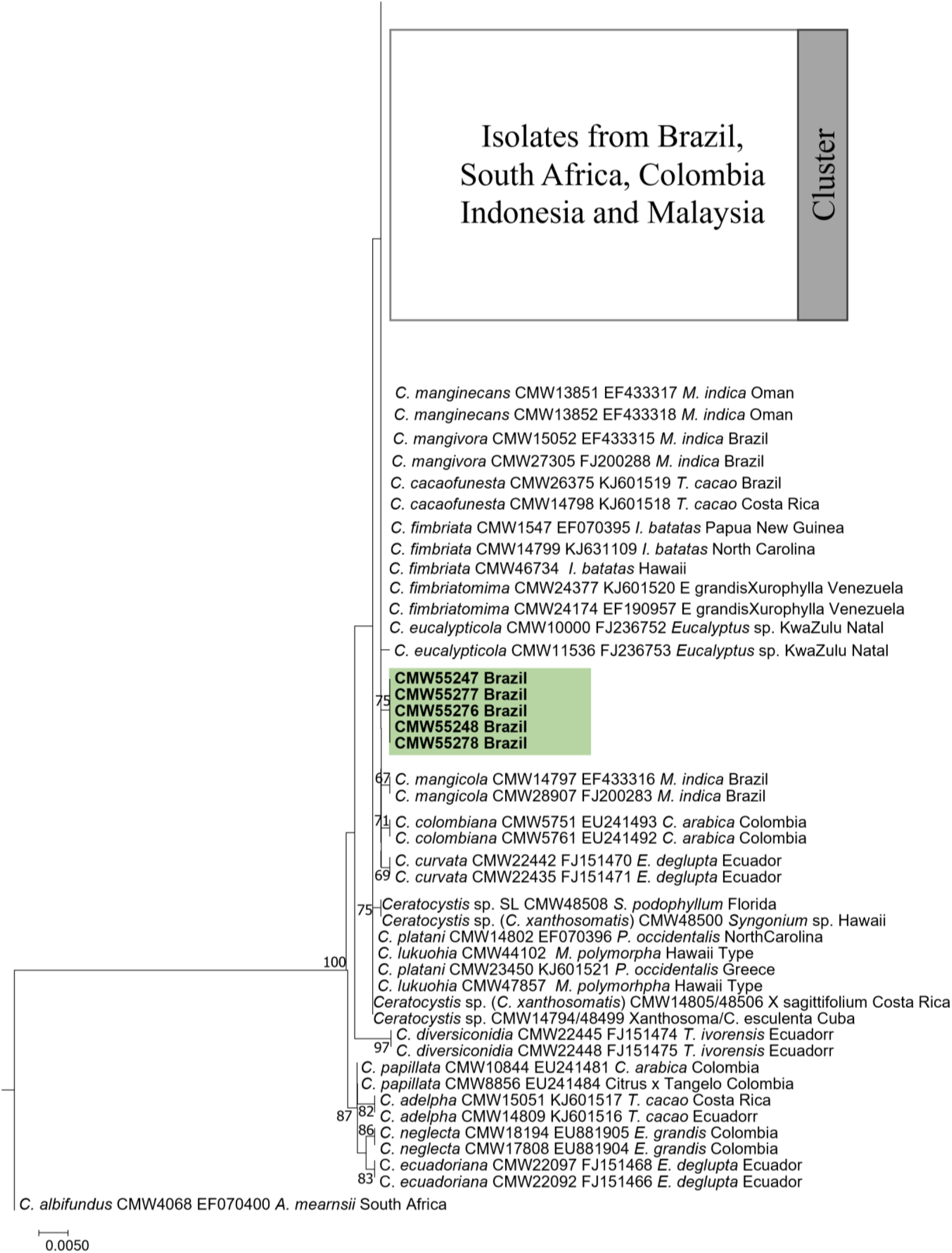
Phylogenetic tree based on maximum likelihood (ML) analysis of tef1 sequences for *Ceratocystis* species in the LAC and *Ceratocystis* isolates used in this study (only representative haplotypes per country and host were included in the analysis). Coloured boxes indicate representatives of isolates sequenced in this study from five regions (Brazil, Colombia, South Africa, Malaysia, and Indonesia). Isolates sequences in this study clustered together with Lineage 1 (*C. eucalypticola*), Lineage 2 (*C. manginecans*), isolates previously designated as *C. mangicola*, *C. mangivora* and the ex-type isolates of the species *C. fimbriatomima, C. cacaofunesta* and *C. fimbriata*. Several isolates from Brazil formed a statistically supported independent monophyletic clade and are highlighted in green. Bootstrap values above 50% are shown. For details regarding specific isolates please refer to the Supplementary Table 1.

**Supplementary Fig. 4.**
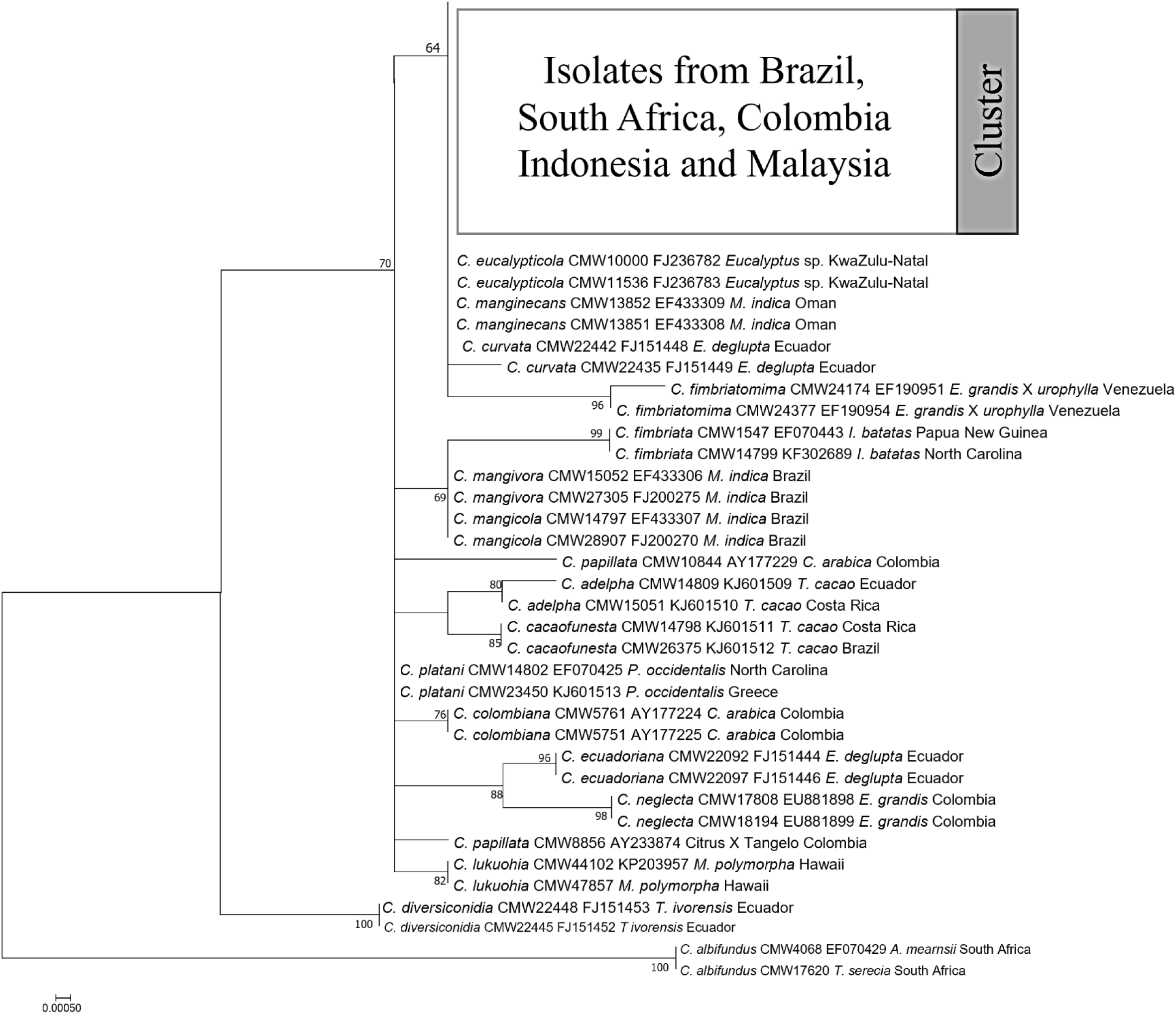
Phylogenetic tree based on maximum likelihood (ML) analysis of βT 1 sequences for *Ceratocystis* species in the LAC and *Ceratocystis* isolates used in this study (only representative haplotypes per country and host were included in the analysis). Coloured boxes indicate representatives of isolates sequenced in this study from five regions (Brazil, Colombia, South Africa, Malaysia, and Indonesia). Isolates sequences in this study clustered together with Lineage 1, Linegae 2 and *C. curvata*. Bootstrap values above 50% are shown. For details regarding specific isolates please refer to the Supplementary Table 1.

**Supplementary Fig. 5.**
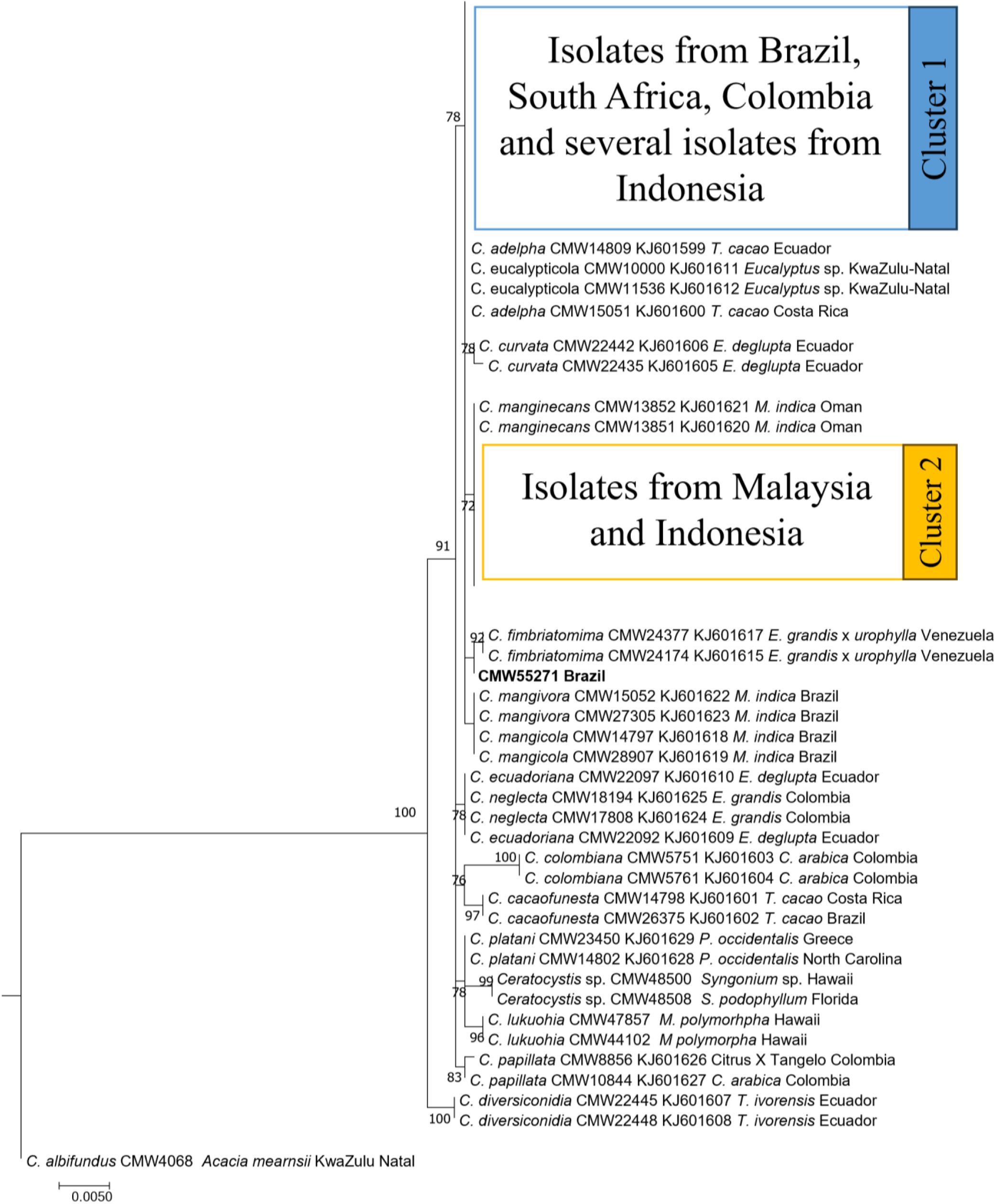
Phylogenetic tree based on maximum likelihood (ML) analysis of rpb2 sequences for *Ceratocystis* species in the LAC and *Ceratocystis* isolates used in this study (only representative haplotypes per country and host were included in the analysis). Coloured boxes indicate representatives of isolates sequenced in this study from five regions (Brazil, Colombia, South Africa, Malaysia, and Indonesia). Isolates from Brazil, South Africa, Colombia, and several isolates from Indonesia screened in this study cluster with Lineage 1 and *C. aldepha* (blue). Isolates from Malaysia and Indonesia formed a statistically supported monophyletic clade with Lineage 2 (yellow). Bootstrap values above 50% are shown. For details regarding specific isolates please refer to the Supplementary Table 1.

**Supplementary Fig. 6.**
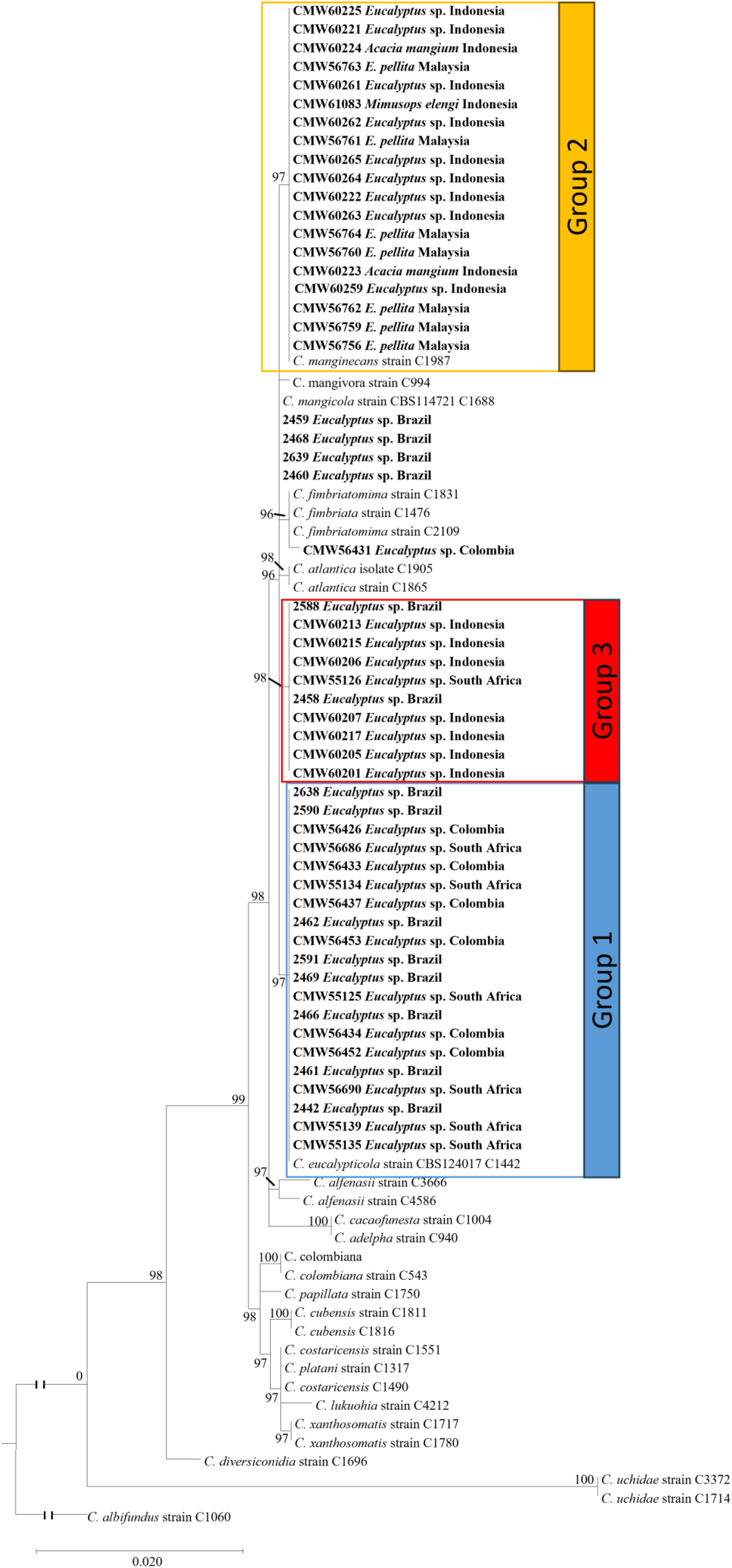
Phylogenetic tree based on maximum likelihood (ML) analysis of MAT1 sequences for *Ceratocystis* species in the LAC and *Ceratocystis* isolates used in this study. Isolates in bold and highlighted in coloured blocks are the isolates sequenced in this study. Isolates in the yellow block formed a statistically supported monophyletic clade with Lineage 2 and consisted of isolates predominantly from Malaysia and Indonesia. Isolates in the blue block formed a statistically supported monophyletic clade with Lineage 1 and consisted of isolates predominantly from Brazil, South Africa, Colombia, and Indonesia. Isolates in the red block formed a statistically supported independent monophyletic clade and consisted of isolates predominantly from Brazil. The remaining isolates grouped with the ex-type isolate previously designated as *C. mangicola* and consisted of isolates from Brazil and a single isolate from Colombia clustered on its own but was closely related to *C. fimbriatomima.* Bootstrap values above 50% are shown.

**Supplementary Fig. 7.**
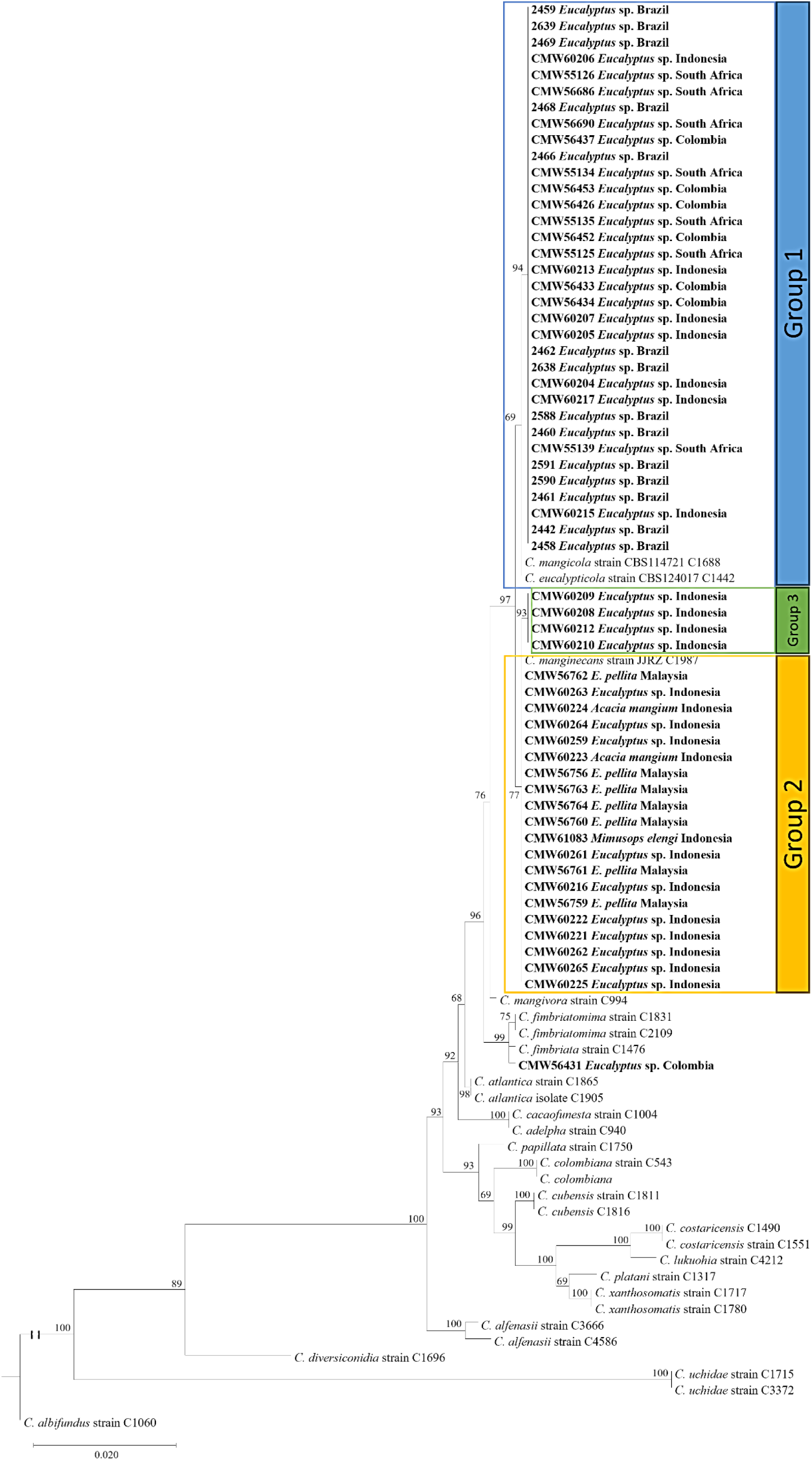
Phylogenetic tree based on maximum likelihood (ML) analysis of MAT2 sequences for *Ceratocystis* species in the LAC and *Ceratocystis* isolates used in this study. Isolates in bold and highlighted in coloured blocks are the isolates sequenced in this study. Most isolates, excluding those from Malaysia, grouped together forming a unique monophyletic clade with high statistical support (blue block). Isolates from Malaysia and Indonesia formed a statistically supported monophyletic clade with Lineage 2 (yellow block). Four isolates from Indonesia formed a unique monophyletic clade with high statistical support (Green block) and a single isolate from Colombia clustered on its own but was closely related to *C. fimbriata* strain C1476 *and C. fimbriatomima*.

**Supplementary Fig. 8.**
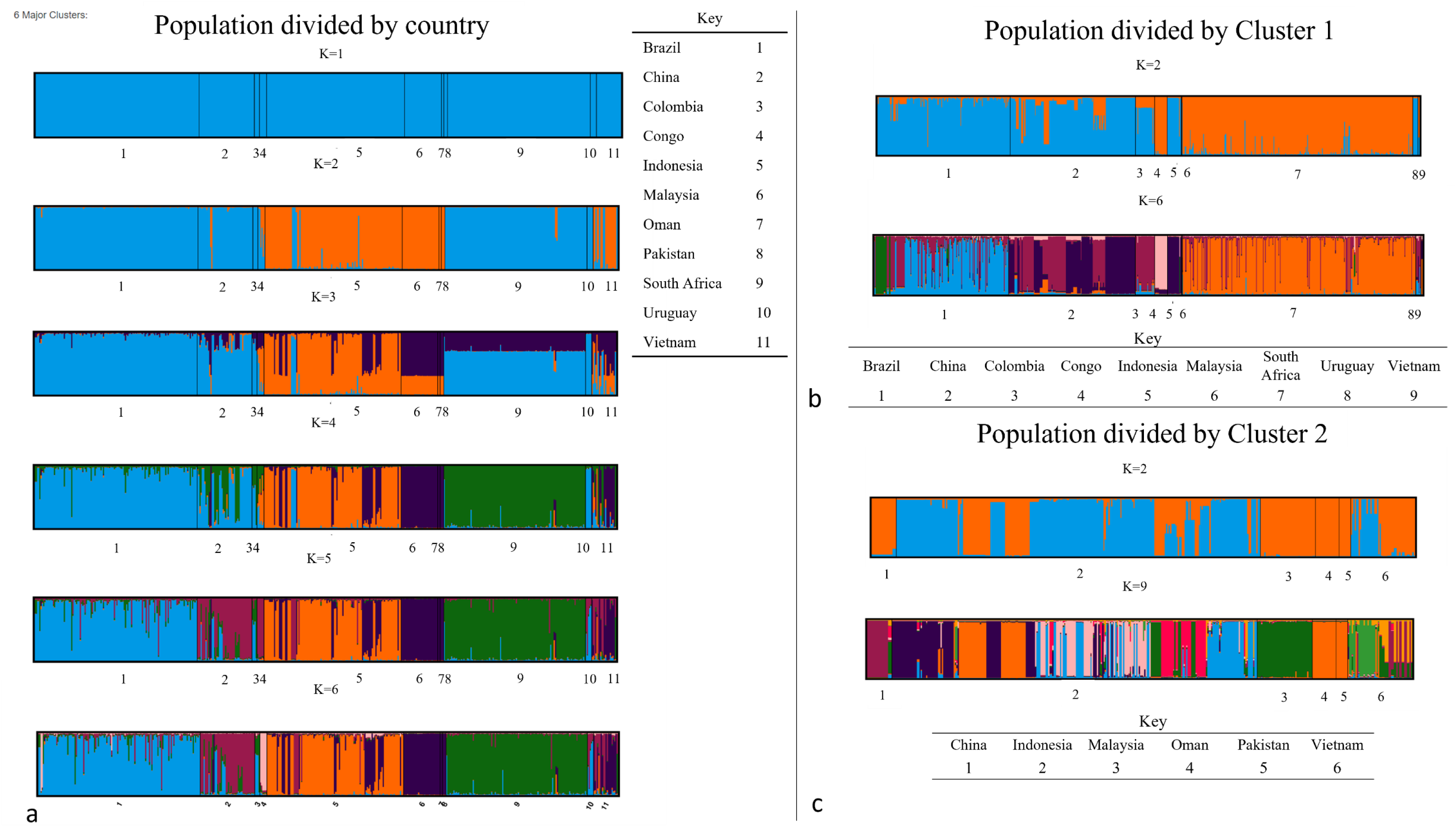
Analysis of population structure (K) of a dataset of the *Ceratocystis* isolates. a) Bar graphs of all possible clusters (K=1–6) for clone-corrected dataset of entire population divided by country. b) Subsequent STRUCTURE analyses conducted on isolates from Cluster 1. The optimal K values of 2 and 6 were suggested for the non-clone-corrected dataset of isolates from Cluster 1 divided by country. c) Subsequent STRUCTURE analyses conducted on isolates from Cluster 2. The optimal K values of 2 and 9 were suggested for the non-clone-corrected dataset of isolates from Cluster 2 divided by country. Each individual is represented by a single vertical line and the colours indicate the relatedness of an isolate to a specific cluster.

**Supplementary Fig. 9.**
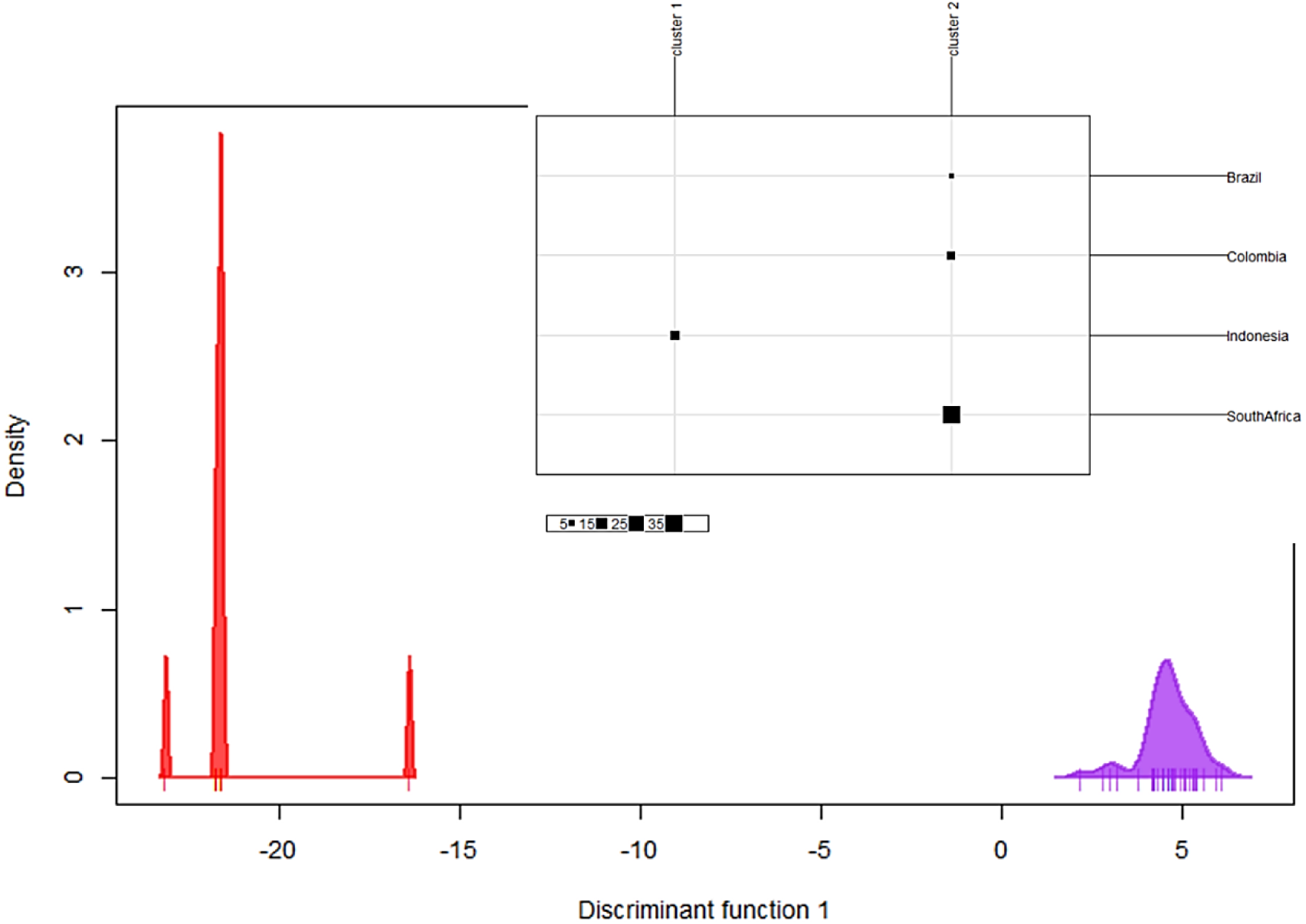
Discriminant Analysis of Principal Components (DAPC) of genetic clusters for the isolates that indicated a mixed ancestry. DAPC plot showing the distribution of isolates in two distinct bell curves. Individuals from Indonesia were exclusive to group 1 and individuals from Brazil, Colombia and South Africa were exclusive to group 2.

**Supplementary Fig. 10.**
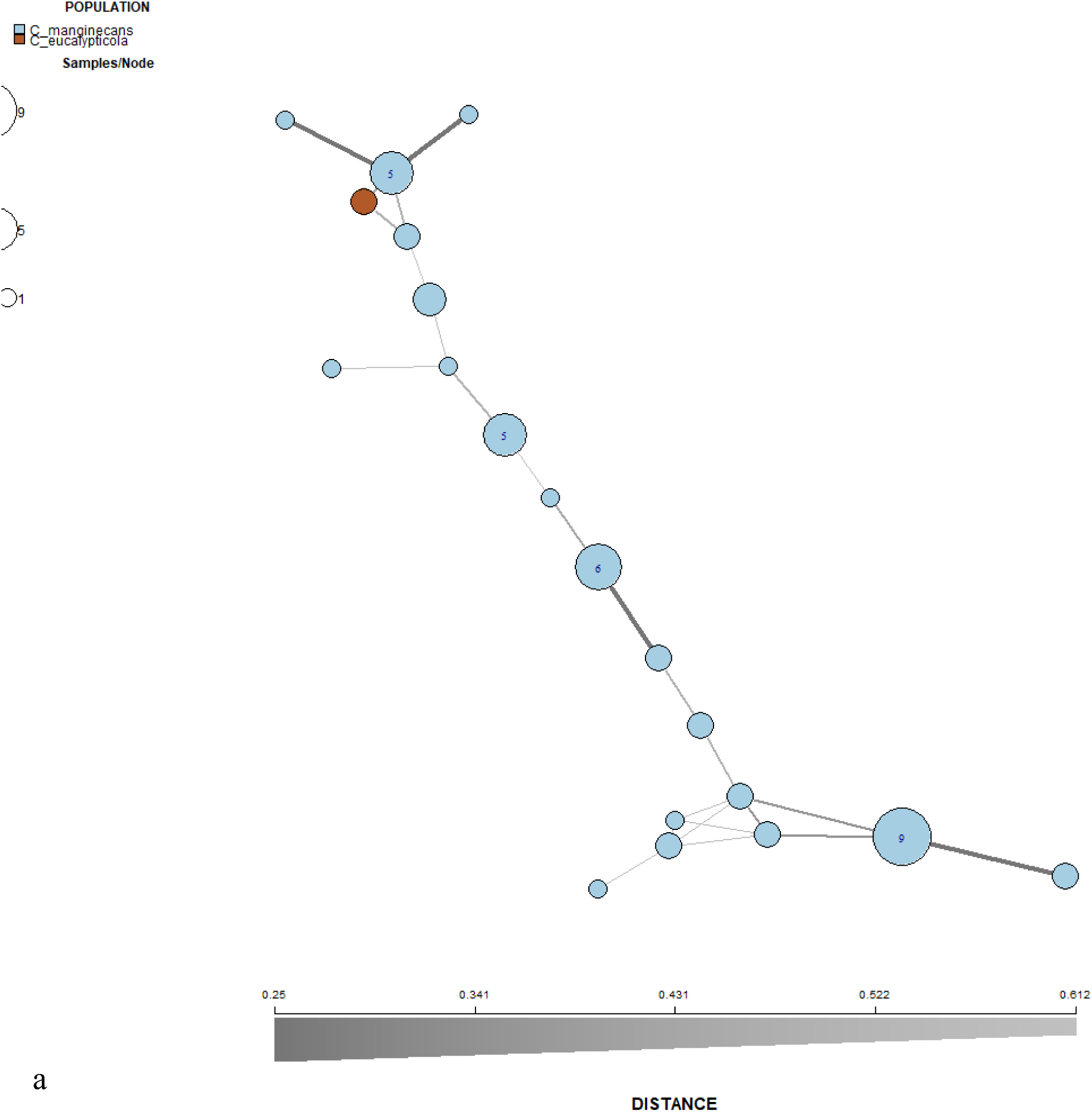
Minimum spanning networks (MSN) showing the relationship of *Ceratocystis* isolates in each country based on Edwards genetic distance. Each node represents one multilocus genotype (MLG) and the size of the node is proportional to the number of individuals with that MLG. The colour gradient from dark to light represents the degree of divergence, where dark lines denote closer genetic similarity and light lines denote greater divergence. 10 a. Minimum spanning networks (MSN) showing the relationship of *Ceratocystis* isolates in Vietnam. Nodes are coloured according to clusters. Brown indicates isolates in cluster 1 and blue indicates cluster 2.

**Supplementary Fig. 10b.**
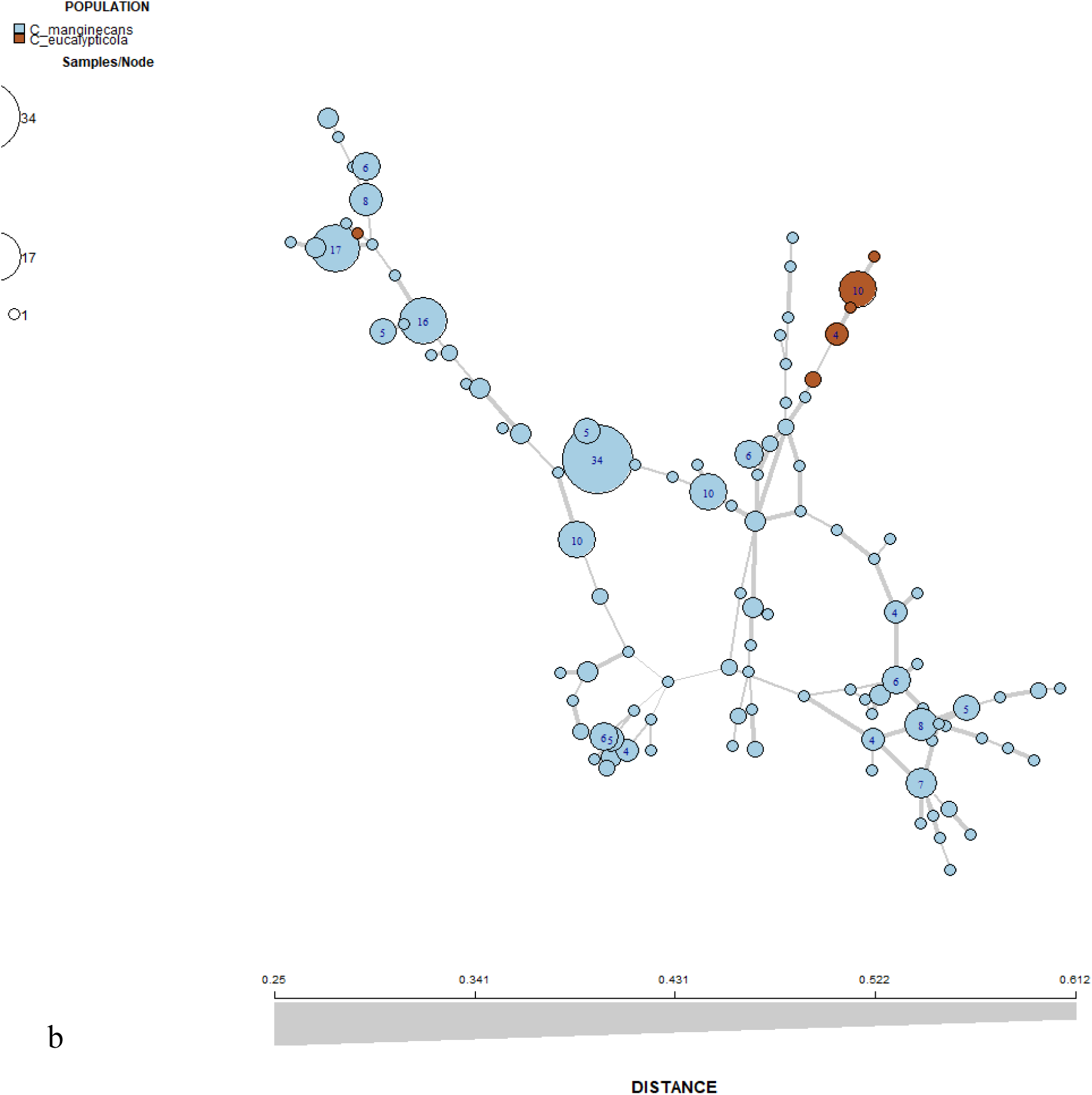
Minimum spanning networks (MSN) showing the relationship of *Ceratocystis* isolates in Indonesia. Nodes are coloured according to clusters. Brown indicates isolates in cluster 1 and blue indicates cluster 2.

**Supplementary Fig. 10c.**
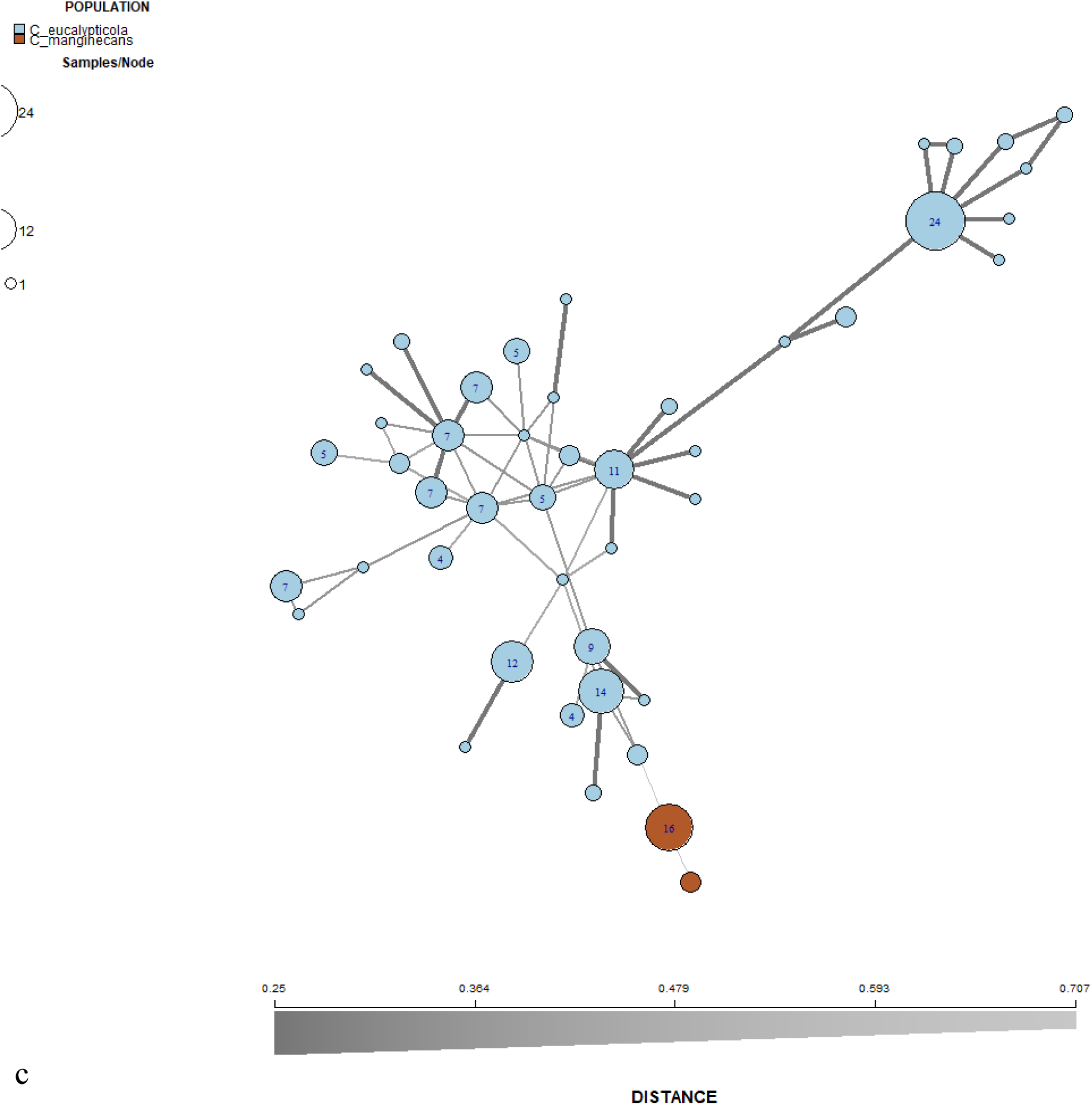
Minimum spanning networks (MSN) showing the relationship of *Ceratocystis* isolates in China. Nodes are coloured according to clusters. Brown indicates isolates in cluster 1 and blue indicates cluster 2.

**Supplementary Fig. 10d.**
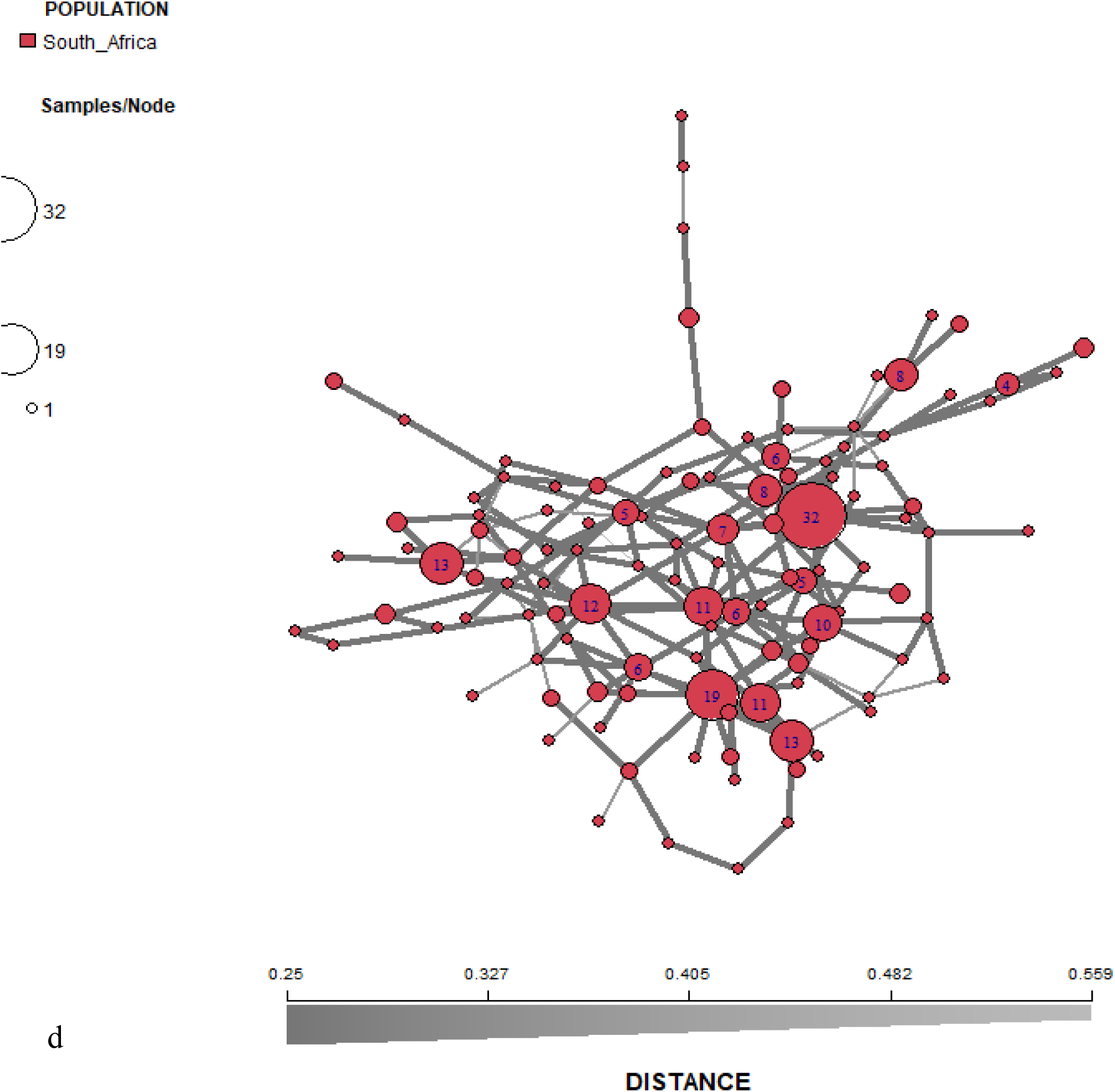
Minimum spanning networks (MSN) showing the relationship of *Ceratocystis* isolates in South Africa.

**Supplementary Fig. 10e.**
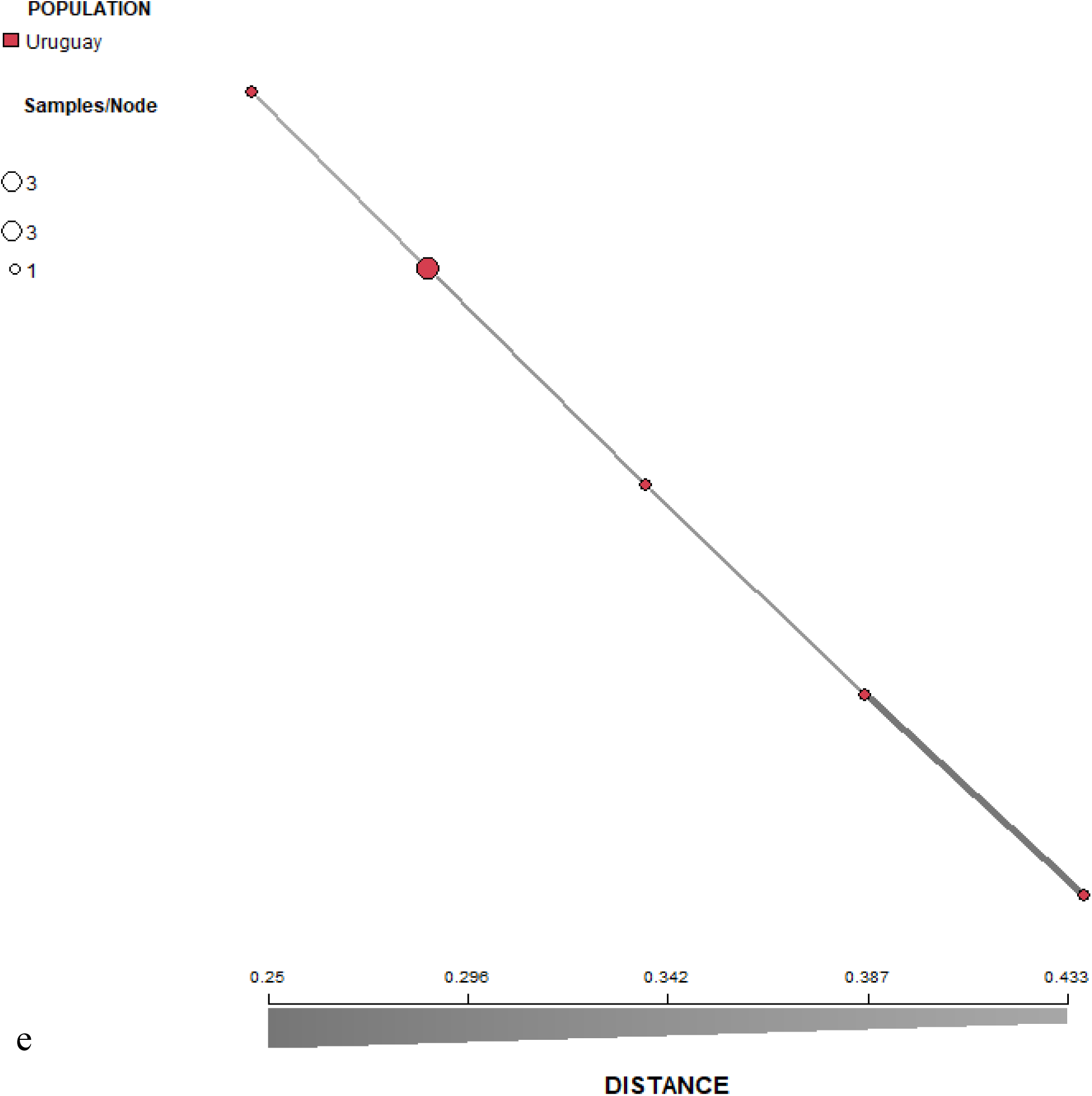
Minimum spanning networks (MSN) showing the relationship of *Ceratocystis* isolates in Uruguay.

**Supplementary Fig. 10f.**
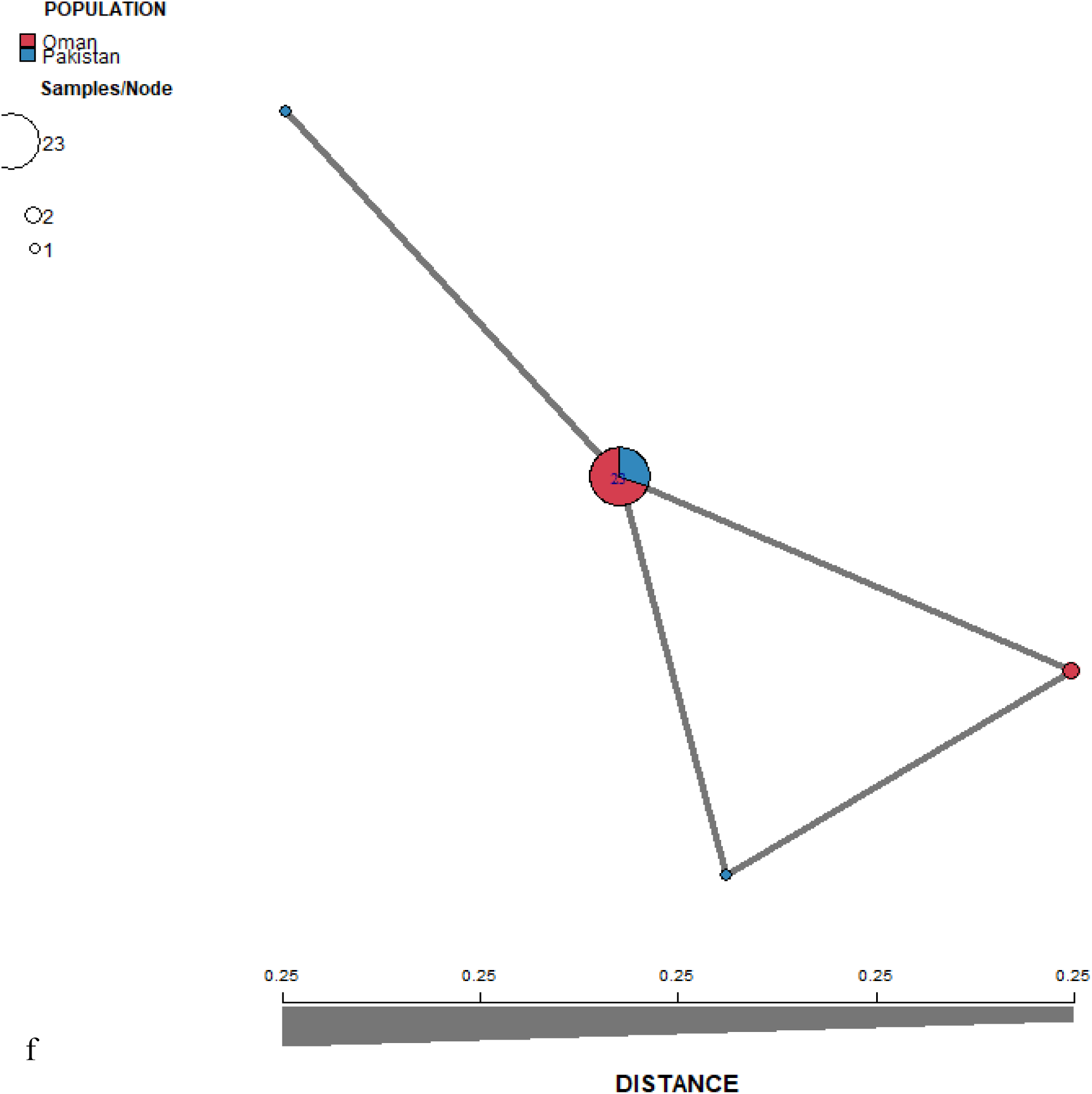
Minimum spanning networks (MSN) showing the relationship of *Ceratocystis* isolates in Oman and Pakistan.

**Supplementary Fig. 10g.**
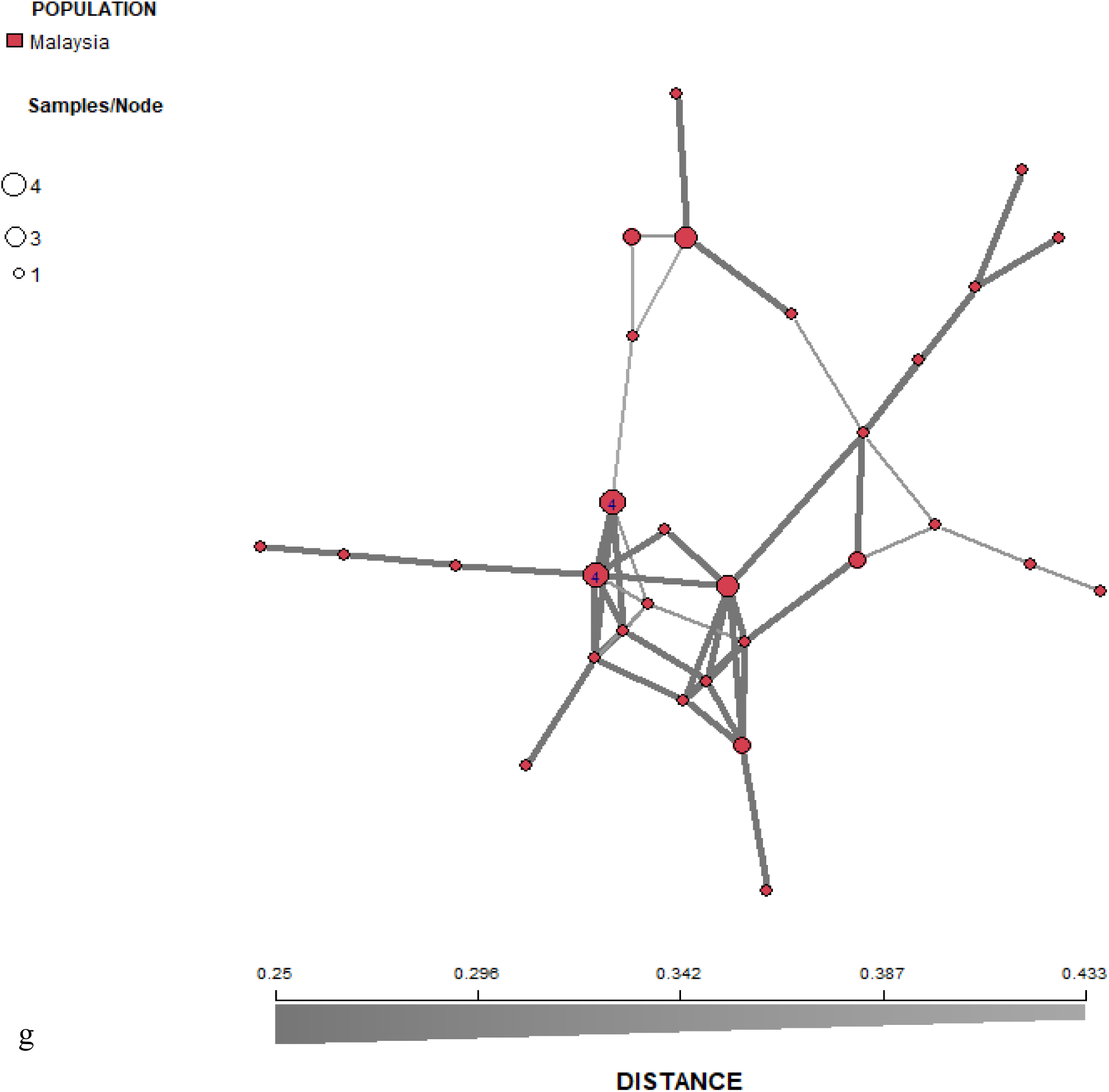
Minimum spanning networks (MSN) showing the relationship of *Ceratocystis* isolates in Malaysia.

**Supplementary Fig. 10h.**
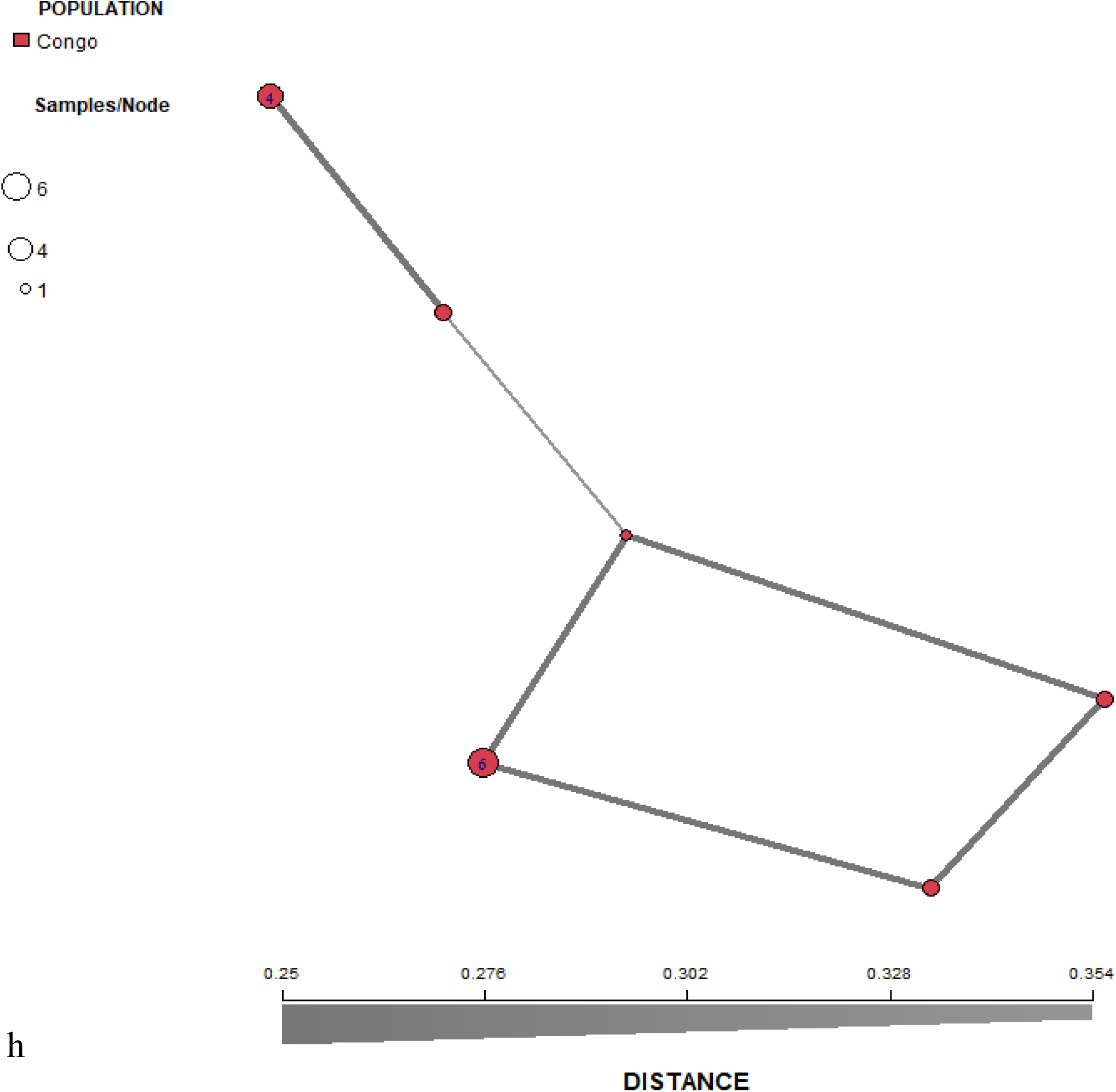
Minimum spanning networks (MSN) showing the relationship of *Ceratocystis* isolates in Congo.

**Supplementary Fig. 10i.**
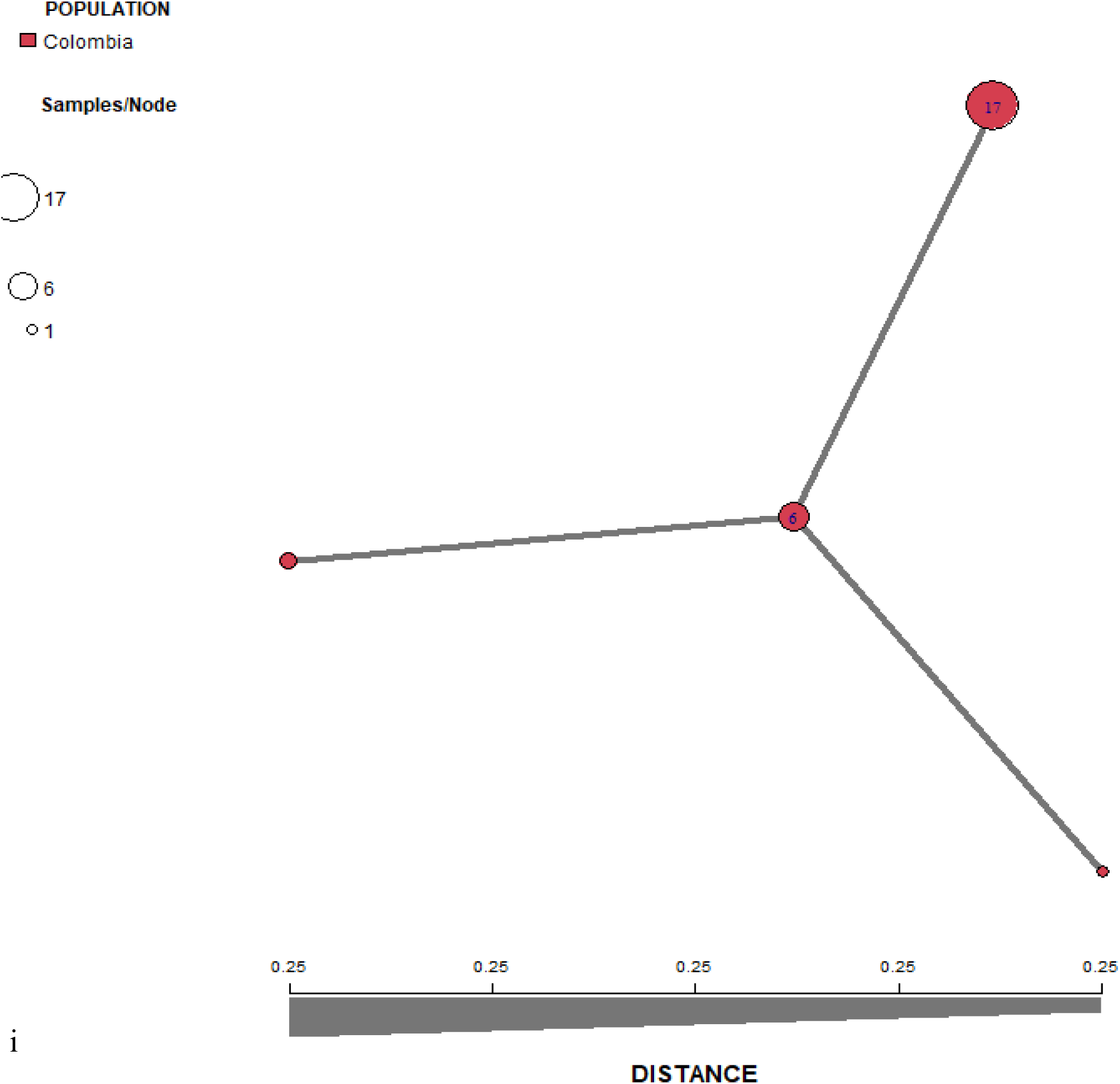
Minimum spanning networks (MSN) showing the relationship of *Ceratocystis* isolates in Colombia.

**Supplementary Fig. 10f.**
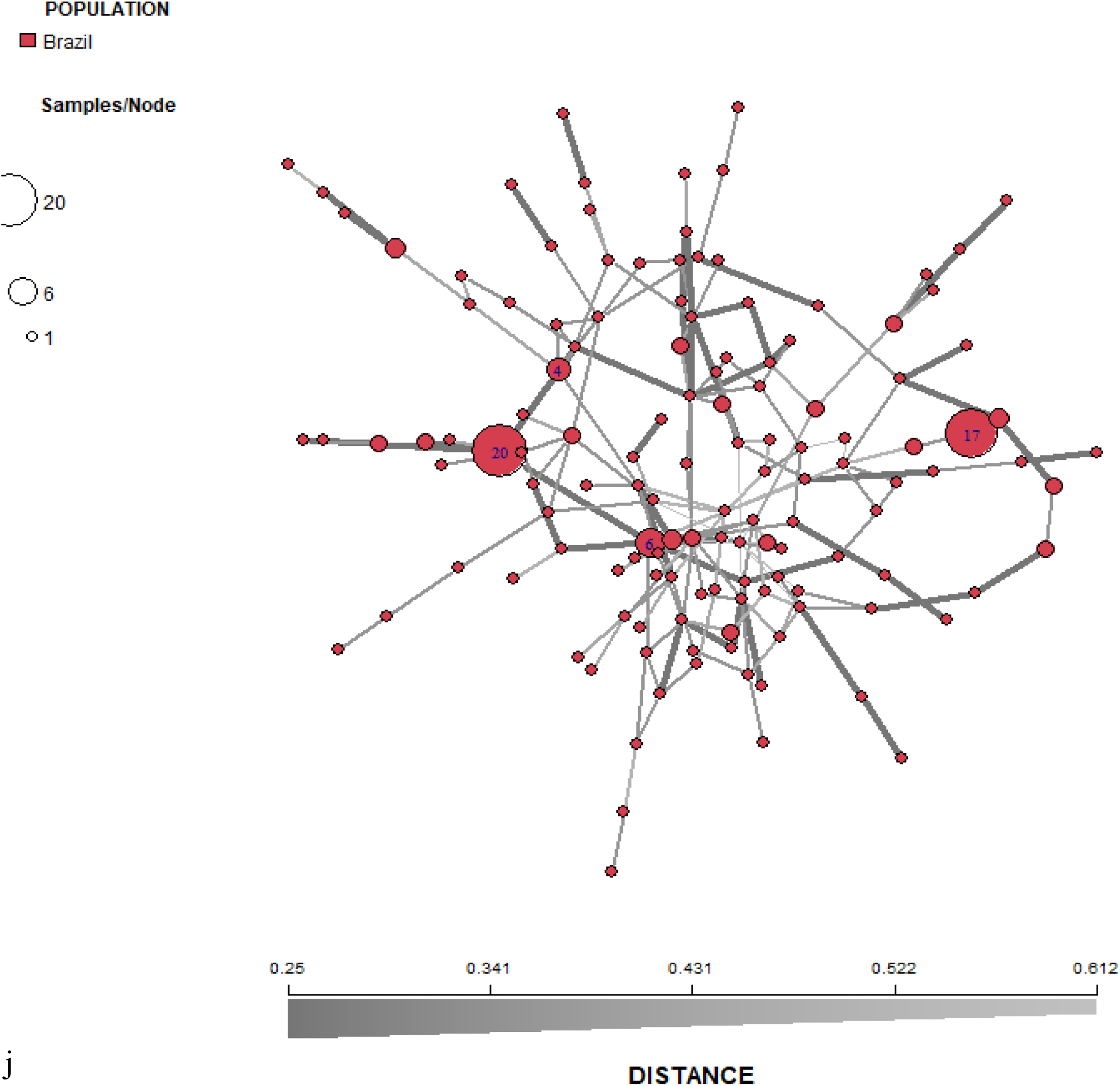
Minimum spanning networks (MSN) showing the relationship of *Ceratocystis* isolates in Brazil.

**Supplementary Fig. 11.**
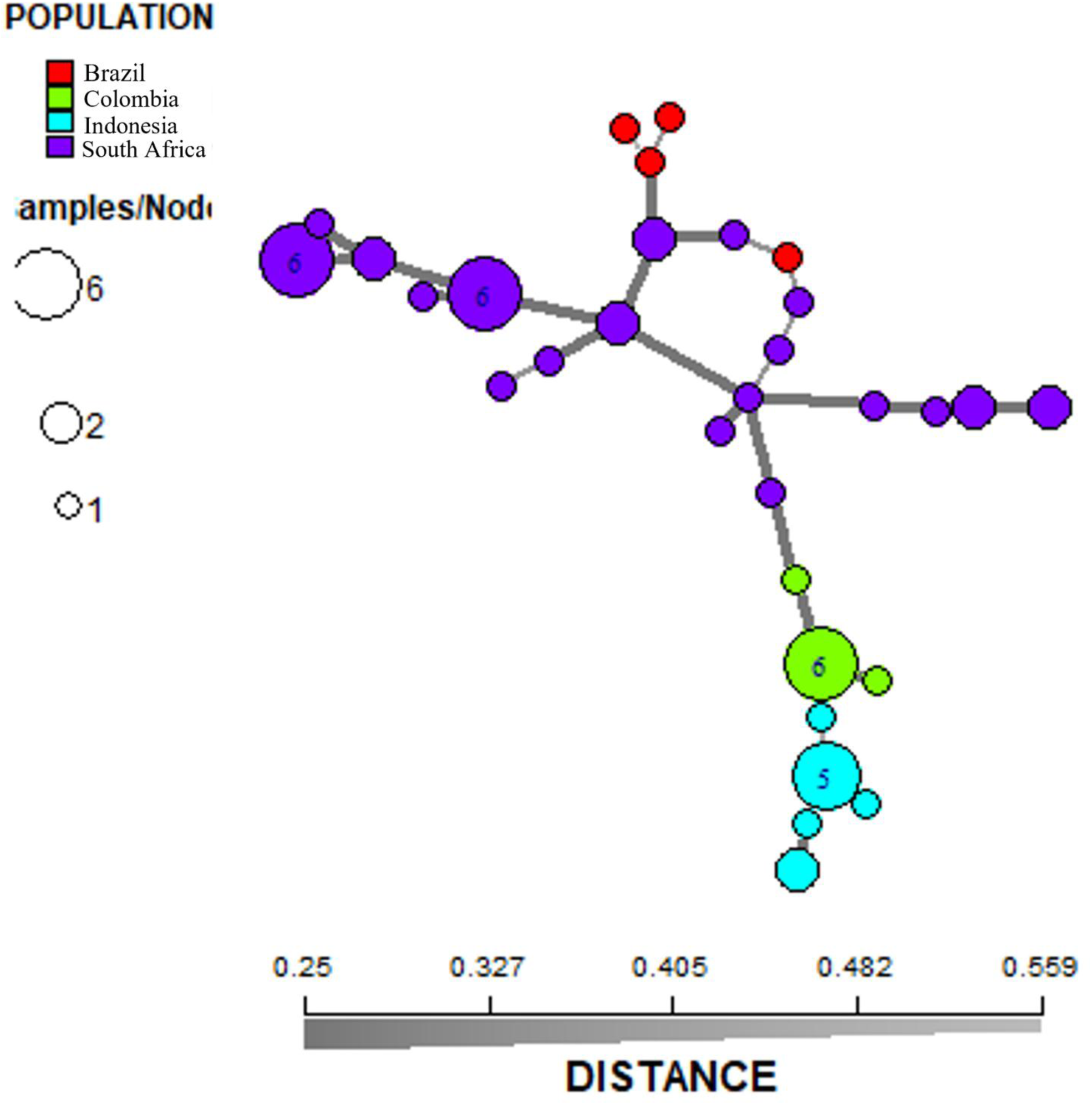
Minimum spanning networks (MSN) showing the relationship of *Ceratocystis* isolates that indicated a mixed ancestry based on Edwards genetic distance. Each node represents one multilogues genotype (MLG) and the size of the node is proportional to the number of individuals with that MLG. Nodes are coloured according to sampling location. The colour gradient from dark to light represents the degree of divergence, where dark lines denote closer genetic similarity and light lines denote greater divergence.

